# Ion channel expression and electrophysiology of singular human (primary and induced pluripotent stem cell derived) cardiomyocytes

**DOI:** 10.1101/2021.03.04.433834

**Authors:** Christina Schmid, Najah Abi-Gerges, Dietmar Zellner, Georg Rast

## Abstract

Human induced pluripotent stem cell-derived cardiomyocytes (hiPSC-CMs) and primary human cardiomyocytes are used for *in vitro* cardiac safety testing. hiPSC-CMs have been associated with a vast heterogeneity regarding single-cell morphology, beating behavior and action potential duration, prompting a systematic analysis of single-cell characteristics. Previously published hiPSC-CM studies revealed action potentials with nodal-, atrial- or ventricular-like morphology, although ion channel expression of singular hiPSC-CMs is not fully understood. Other studies used single-cell RNA-sequencing, however, these studies did not extensively focus on expression patterns of cardiac ion channels or failed to detect ion channel transcripts. Thus, the current study used a single-cell patch-clamp-RT-qPCR approach to get insights into single-cell electrophysiology (capacitance, action potential duration at 90% of repolarization, upstroke velocity, spontaneous beat rate, and sodium-driven fast inward current) and ion channel expression (HCN4, CACNA1G, CACNA1D, KCNA5, KCNJ4, SCN5A, KCNJ2, CACNA1D, and KCNH2), the combination of both within individual cells, and their correlations in single cardiomyocytes. We used commercially available hiPSC-CMs (iCell cardiomyocytes, atrial and ventricular Pluricytes) and primary human adult atrial and ventricular cardiomyocytes. Recordings of electrophysiological parameters revealed differences between the cell groups and variation within the hiPSC-CMs groups as well as within primary ventricular cardiomyocytes. Expression analysis on mRNA level showed no-clear-cut discrimination between primary cardiac subtypes and revealed both similarities and differences between all cell groups. Higher expression of atrial-associated ion channels in primary atrial cardiomyocytes and atrial Pluricytes compared to their ventricular counterpart indicates a successful chamber-specific hiPSC differentiation. Interpretation of correlations between the single-cell parameters was challenging, as the total data set is complex, particularly for parameters depending on multiple processes, like the spontaneous beat rate. Yet, for example, expression of SCN5A correlated well with the fast inward current amplitude for all three hiPSC-CM groups. To further enhance our understanding of the physiology and composition of the investigated hiPSC-CMs, we compared beating and non-beating cells and assessed distributions of single-cell data. Investigating the single-cell phenotypes of hiPSC-CMs revealed a combination of attributes which may be interpreted as a mixture of traits of different adult cardiac cell types: (i) nodal-related pacemaking attributes are spontaneous generation of action potentials and high HCN4 expression; and (ii) non-nodal attributes: cells have a prominent I_Na_-driven fast inward current, a fast upstroke velocity and a high expression of SCN5A. In conclusion, the combination of nodal- and non-nodal attributes in single hiPSC-CMs may hamper the interpretation of drug effects on complex electrophysiological parameters like beat rate and action potential duration. However, the proven expression of specific ion channels enables the evaluation of drug effects on ionic currents in a more realistic environment than in recombinant systems.

## 1 Introduction

Primary human cardiomyocytes as well as human induced pluripotent stem cell derived cardiomyocytes (hiPSC-CMs) are used as model systems in pharmacological and toxicological drug testing (Magdy, Schuldt, Wu, Bernstein, & Burridge, 2018; Nguyen et al., 2017; Pang et al., 2019; Yoshida & Yamanaka, 2017). They are, for example, recommended in the context of the Comprehensive *in vitro* Proarrhythmia Assay (CiPA) initiative (Wallis et al., 2018). For the assessment of potential acute, electrophysiological side effects of drug candidates, it is essential to know the electrophysiological characteristics and ion channel expression of a cardiomyocyte model. For hiPSC-CMs, expression of cardiac ion channels was reported on a population level and single-cells were shown to display ionic currents (Barbuti, Benzoni, Campostrini, & Dell’Era, 2016; Karakikes, Ameen, Termglinchan, & Wu, 2015; Ma et al., 2011). However, these studies used different subsets of cells for the investigation of bulk ion channel expression and single-cell electrophysiology (e.g. ionic current and action potential measurements).

Working with singularized hiPSC-CMs, we observed heterogeneous cell sizes and morphologies of individual cells. Moreover, we recognized that some cells were contracting while others wereńt and that action potential durations varied immensely between different cells. These first observations of obvious differences between individual cells prompted us to perform a systematic characterization of hiPSC-CMs on the single-cell level and to compare them with primary human cardiomyocytes isolated from donor hearts.

It is well established that cardiomyocytes from different regions of the heart (nodal, atrial and ventricular cardiomyocytes) have different ion channel expression and action potential morphology (Kane & Terracciano, 2017). Heterogeneity of hiPSC-CMs has been mainly assessed with two approaches. On the one hand, single-cell RNA-sequencing was performed. The majority of studies using this approach did not detect distinct transcriptomic cardiac subtype related clusters in hiPSC-CMs (Biendarra-Tiegs, Li, et al., 2019; Friedman et al., 2018; Schmid, Wohnhaas, Hildebrandt, Baum, & Rast, 2020). In contrast, one single-cell RNA-sequencing study reported atrial and ventricular related gene expression (Churko et al., 2018). On the other hand, a second approach used action potential morphology to classify hiPSC-CMs into nodal-, atrial-, and ventricular-like cells (Ma et al., 2011; Yonemizu et al., 2019).

In our opinion, action potential morphology alone is not sufficient to classify hiPSC-CMs into primary subtypes due to missing information about the underlying ionic currents and channels. We complemented the single-cell characterization of human cardiomyocytes with the combination of electrophysiological recordings and ion channel expression in the same cell using a single-cell patch-clamp-RT-qPCR technique. For this purpose, we used three different commercially available hiPSC-CMs (iCell cardiomyocytes, ventricular Pluricytes and atrial Pluricytes) and compared them with primary adult human atrial and ventricular cardiomyocytes. We chose to focus on nine cardiac ion channel genes (HCN4, CACNA1G, CACNA1D, KCNA5, KCNJ4, SCN5A, KCNJ2, CACNA1C and KCNH2) because of their association with recorded electrophysiological parameters and/or their association with primary cardiac subtypes. For the understanding of the present study, it is essential to know that the SCN5A gene (coding for the NaV1.5 channel subunit and conducting I_Na_) is associated with the fast inward current and the upstroke of the action potential in primary atrial and ventricular cardiomyocytes, while it shows no/low expression in primary nodal cells (Kane & Terracciano, 2017; Priest & McDermott, 2015). In primary nodal cardiomyocytes, depolarization is mainly driven by HCN4 (conducting the If current), CACNA1G (coding for CaV3.1 which conducts ICaT) and CACNA1D (coding for CaV1.3 which conducts I_CaL_) and these ion channels are known to contribute to pacemaking in these cells (Baig et al., 2011; Bartos, Grandi, & Ripplinger, 2015; Mesirca, Torrente, & Mangoni, 2014). While these three channels also show expression in atrial cells, no/low expression was detected in ventricular cardiomyocytes (Kane & Terracciano, 2017). The absence of the IK1 current (KCNJ2 and KCNJ4, coding for Kir2.1 and Kir2.3, respectively) is also assumed to contribute to pacemaking in primary nodal cells (Goversen, van der Heyden, van Veen, & de Boer, 2018; Verkerk et al., 2007). The expression of KCNA5 (coding for K_V_1.5 which conducts I_Kur_) is highly associated with primary atrial cardiomyocytes (Kane & Terracciano, 2017) and I_CaL_ (CACNA1C coding for CaV1.2) and I_Kr_ (KCNH2 coding for K_V_11.1/HERG) are related to action potential duration (Rast, Kraushaar, Buckenmaier, Ittrich, & Guth, 2016). First, we compared the electrophysiological and transcriptomic results within and between the groups on a population and a single-cell level. Next, we assessed whether beating and non-beating cells within a hiPSC-CM group differ with respect to electrophysiology or ion channel expression. In addition, we investigated whether distributions of single-cell parameters help to determine subpopulations or whether single-cell hiPSC-CM phenotypes (the combination of electrophysiological and transcriptomic attributes) could be classified into primary cardiac subtypes (nodal, atrial, and ventricular). Finally, we investigated the single-cell relationship between ion channel transcripts and electrophysiological parameters.

## 2 Material and Methods

### 2.1 Culture of hiPSC-derived cardiomyocytes

Three different commercially available hiPSC-derived cardiomyocyte products were purchased for this study: iCell cardiomyocytes (Fujifilm Cellular Dynamics, Madison, WI, USA), ventricular Pluricytes and custom-made atrial Pluricytes (Ncardia, Gosselies, Charleroi, Belgium). iCell cardiomyocytes originated from 5 different vials of the same lot, ventricular Pluricytes from 3 vials of the same lot and atrial Pluricytes from 2 vials of the same lot.

#### 2.1.1 iCell cardiomyocytes

iCell cardiomyocytes were cultured as described previously (Schmid et al., 2020). Briefly, the following steps were performed: After thawing, iCell cardiomyocytes (product number 01434, Fujifilm Cellular Dynamics, Madison, WI, USA) were centrifuged (180 x g for 5 min) and resuspended in Plating Medium (Fujifilm Cellular Dynamics, Madison, WI, USA). Cells were seeded into fibronectin-coated (Sigma-Aldrich, St. Louis, MO, USA) 24-well plates (#142485, Thermo Fisher Scientific, Waltham, MA, USA) at a density of 1.3 x 10^5^ cells/well, followed by incubation at 37°C in ambient atmosphere supplemented with 7.5% CO_2_ and 90% relative humidity. Plating Medium was switched to Maintenance Medium (Fujifilm Cellular Dynamics, Madison, WI, USA) after 48 h. Subsequently, the Maintenance Medium was replaced every 3^rd^ to 4^th^ day. Formation of a monolayer and synchronous beating was observed after 3-5 days. iCell cardiomyocytes were singularized for experiments 12-39 days after thawing.

#### 2.1.2 Ventricular and atrial Pluricytes

Ventricular (product number PCK-1.5, Ncardia, Gosselies, Charleroi, Belgium) and atrial Pluricytes (custom production, Ncardia, Gosselies, Charleroi, Belgium) were thawed and cultured according to manufacturer’s instructions. Briefly, Pluricytes were thawed, subsequently centrifuged (250 x g, 3 min) and resuspended in Pluricyte Cardiomyocyte Medium (Ncardia, Gosselies, Charleroi, Belgium). Cells were seeded into fibronectin-coated (Sigma-Aldrich, St. Louis, MO, USA) 24-well plates (#142485, Thermo Fisher Scientific, Waltham, MA, USA) at a density of 1-2 x 10^5^ cells/well. Pluricytes were then incubated at 37°C in ambient atmosphere supplemented with 5% CO_2_ and 90% relative humidity. The first medium change was performed after 24 h. Subsequently, medium changes were performed every 2^nd^ to 3^rd^ day. After 3-5 days, Pluricytes formed a monolayer and started to beat synchronously. On day 7-20 (atrial) or day 13-23 (ventricular) after thawing, Pluricytes were singularized for experiments.

### 2.2 Singularizing hiPSC-derived cardiomyocytes

To investigate single cells, singularization of monolayer hiPSC-CM cultures was performed by splitting confluent cells of a single well into 4 - 6 new wells of a 24-well plate: cardiomyocyte monolayers were washed with D-PBS. iCell cardiomyocytes were then exposed to 0.25% trypsin-EDTA (Sigma-Aldrich, St. Louis, MO, USA) for 8 min at 37°C. Ventricular and atrial Pluricytes were exposed to TrypLE Express (Thermo Fisher Scientific, Waltham, MA, USA) for 10 min at 37°C. Afterwards, Maintenance Medium (Fujifilm Cellular Dynamics, Madison, WI, USA, for iCells) or Pluricyte Cardiomyocytes Medium (Ncardia, Gosselies, Charleroi, Belgium, for Pluricytes) was added to the cardiomyocytes, and pipetted up and down ten times, using a P1000 pipette (Eppendorf AG, Hamburg, Germany). Then, the cell suspension was transferred into a 50 mL-tube (Corning, New York, NY, USA) and diluted with Maintenance Medium or Pluricyte Cardiomyocyte Medium up to a minimum of 15 mL in total. iCell cardiomyocyte suspensions were centrifuged at 180 x g for 5 min. Pluricyte suspensions were centrifuged at 125 x g for 5 min. Supernatant was aspirated and remaining cells were resuspended in an adequate volume of Maintenance Medium or Pluricyte Cardiomyocyte Medium to achieve a 1:4-1:6-splits (seeding volume: 200 µL cell suspension per well). Cells were seeded on Matrigel (human embryonal stem cell (hESC)-qualified, Corning, New York, NY, USA) coated, 12 mm glass coverslips (Thermo Fisher Scientific, Waltham, MA, USA) placed in 24 well plates (#142485, Thermo Fisher Scientific, Waltham, MA, USA) already containing 1 mL of Maintenance Medium or Pluricytes Cardiomyocyte Medium per well. Between day 1 and 3 after singularizing, hiPSC-CMs were used for experiments.

### 2.3 Primary human cardiomyocytes

The procurement of donor hearts and the isolation of primary human cardiomyocytes was performed at AnaBios Corporation (San Diego, CA, USA).

#### 2.3.1 Procurement of donor hearts

All methods were carried out in accordance with relevant guidelines and regulations. All human hearts used for this study were non-transplantable and ethically obtained by legal consent (first person or next-of-kin) from cadaveric organ donors in the United States. Our recovery protocols and *in vitro* experimentation were pre-approved by IRBs (Institutional Review Boards) at transplant centers within the US OPTN (Organ Procurement Transplant Network). Furthermore, all transfers of the donor hearts are fully traceable and periodically reviewed by US Federal authorities. Donor characteristics, heart number and donor identifier are shown in Supp. Table 1 and exclusion criteria were previously described (Page et al., 2016).

**Table 1:**
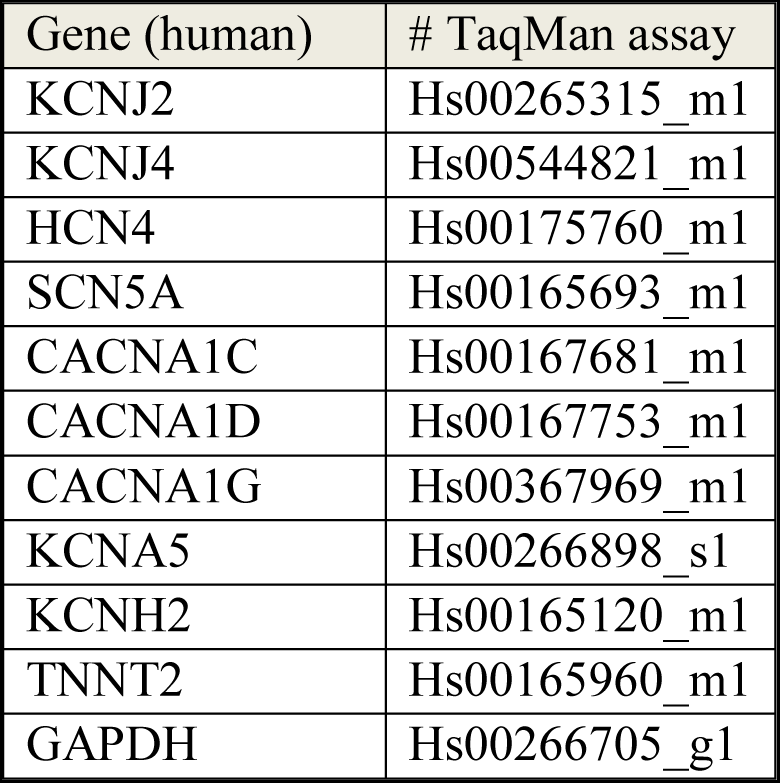
TaqMan gene expression assays used for experiments. Probes span introns whenever possible.

| Gene (human) | # TaqMan assay |
| --- | --- |
| KCNJ2 | Hs00265315_m1 |
| KCNJ4 | Hs00544821_m1 |
| HCN4 | Hs00175760_m1 |
| SCN5A | Hs00165693_m1 |
| CACNA1C | Hs00167681_m1 |
| CACNA1D | Hs00167753_m1 |
| CACNA1G | Hs00367969_m1 |
| KCNA5 | Hs00266898_s1 |
| KCNH2 | Hs00165120_m1 |
| TNNT2 | Hs00165960_m1 |
| GAPDH | Hs00266705_g1 |

#### 2.3.2 Isolation and storage of primary human cardiomyocytes

Upon arrival at the laboratory, hearts were re-perfused with ice-cold proprietary cardioplegic solution and adult human primary ventricular and atrial myocytes were isolated enzymatically from the ventricles or atria, respectively (Nguyen et al., 2017; Page et al., 2016). Single-cell suspensions were stored in proprietary storage solution at 4 °C until use. Primary ventricular cardiomyocytes were used 1 - 5 days after isolation and atrial cardiomyocytes were used 1- 4 days after isolation by trickling cell solution on laminin-coated (24 h at 4 °C or 2 h at rt) glass coverslips and letting them attach for 10 min at rt.

### 2.4 Electrophysiological measurements and cell harvesting

All procedures were performed under ribonuclease (RNase)-free conditions to the largest extent feasible: Dedicated RNase-free workbench, equipment, and material was used, either bought labeled RNase-free or cleaned with RNaseZap (Thermo Fisher Scientific, Waltham, MA, USA) or ElectroZap (Thermo Fisher Scientific, Waltham, MA, USA) or treated with low-pressure plasma for 15 min (MiniFlecto plasmacleaner, Plasma Technology, Herrenberg-Gültstein, Germany). Moreover, RNase-free LoBind tubes (Thermo Fisher Scientific, Waltham, MA, USA), barrier tips (Eppendorf) and UltraPure DNase/RNase-free water (not DEPC treated, Thermo Fisher Scientific, Waltham, MA, USA) were used. Coverslips with singularized cells were placed at the bottom of a bath chamber mounted on an inverted microscope and covered with extracellular (EC-) solution (137.0 mM NaCl, 2.7 mM KCl, 8.0 Na_2_HPO_4_, 2.0 mM KH_2_PO_4_, 0.5 mM MgCl_2_*6 H_2_O, 0.9 mM CaCl2, 5.0 mM glucose). Patch pipettes (made from borosilicate glass tubing (Hilgenberg, Malsfeld, Germany) using a horizontal puller) were filled with intracellular (IC-) solution (For experiments with hiPSC-CMs: 130.0 mM K-aspartate, 5.0 mM MgCl_2_*6 H_2_O, 5.0 mM EGTA, 4.0 mM K_2_ATP, 10.0 mM HEPES, pH 7.2 with KOH. For experiments with primary cardiomyocytes: 100.0 mM K-aspartate, 25.0 mM KCl, 1.0 mM MgCl_2_ *6 H_2_O, 10.0 mM EGTA, 5.0 mM K_2_ATP, 5.0 mM HEPES, pH 7.2 with KOH). Microelectrodes had an open tip resistance between 2.3 and 3.5 MΩ. Membrane currents and action potentials were recorded at 32 °C in the whole cell configuration using an EPC-10 patch clamp amplifier (HEKA Electronics, Lambrecht, Germany) and PatchMaster software (HEKA Electronics, Lambrecht, Germany).

Before starting the electrophysiological measurements, the spontaneous beating frequency of hiPSC-derived cardiomyocytes was counted visually for 30 s. Recordings were started immediately after establishment of the whole-cell configuration, subsequent measurements were performed as fast as possible to avoid both diffusion of mRNA into the pipette and mRNA degradation. To record the fast inward current, cells were clamped at a holding potential of - 100 mV (to make sure sodium channels were not inactivated) followed by a voltage step to -30 mV for 100 ms. Current and voltage signals were Bessel filtered at 10 kHz before being digitized at 20 kHz. Rs was compensated to a level that safely allowed recording devoid of ringing throughout recordings (typically 50%). Inward current peak amplitudes were obtained by subtracting holding current at -100 mV from the inward peak current at -30 mV. Action potential recordings were obtained by a “gentle switch” to current-clamp mode at a holding potential of -80 mV, i.e. the command current in current clamp mode is automatically set to the average value determined in voltage clamp mode for the holding potential. To ensure comparability of action potentials the command current was adjusted to keep cells in a range of -76 mV to -84 when necessary. Action potentials of hiPSC-CMs were triggered by injecting rectangular current pulses (500 – 1000 pA for 2-3 ms) every 3 s. For recordings from primary cardiomyocytes, current pulses were 1000-4000 pA for 1-3 ms every 3 s. Three replicate voltage clamp sweeps and three replicate current clamp sweeps per cell were used for analysis. Current and action potential traces were exported to Microsoft Excel (Redmond, WA, USA) for analysis. The following electrophysiological parameters were determined: Capacitance [pF], beat rate [bpm], inter-beat interval [ms] (calculation from counted beat rate, includes only beating cells, as inter-beat interval would be not defined for rate = 0 but we wanted to investigate the beating cells as a sub-group), fast inward current amplitude [nA], fast inward current density (current normalized to cell capacitance; [pA/pF]), upstroke velocity of action potential (AP) [V/s] and APD90 (AP duration at 90% repolarization, for calculation see equation below) [ms].

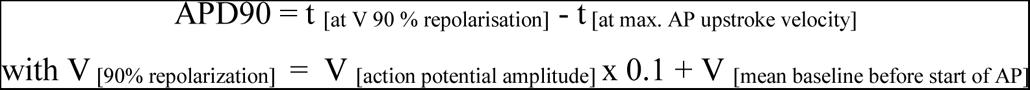

Graphical representations were generated using GraphPad Prism (GraphPad software, San Diego, CA).

After electrophysiological measurements, the micropipette (filled with IC-solution) used for recordings was gently removed from the cell, allowing the cell to re-seal. A large-bore pipette filled with a low amount (approximately 1 µL) of RNase-free phosphate buffered saline (PBS) (Invitrogen, Carlsbad, CA, USA) was used to aspirate the same cell under visual control. After aspiration, the tip of the pipette was broken into a single nuclease-free 0.2 mL PCR-tube (Eppendorf AG, Hamburg, Germany), pre-filled with 9 µL Single Cell Lysis Solution and 1 µL Single Cell DNAse, Thermo Fisher Scientific, Waltham, MA, USA). For the subsequent single-cell RT-qPCR procedure see 2.5. One coverslip with cardiomyocytes was used for a maximum of 30 min. In total, 72 iCells, 24 atrial Pluricytes, 29 ventricular Pluricytes, 40 primary atrial human cardiomyocytes and 90 primary ventricular human cardiomyocytes were collected. It was not possible to determine every parameter for every single cell for technical reasons; therefore, *n* may vary between parameters.

### 2.5 Single-cell RT-qPCR

Single-cell mRNA expression of nine cardiac ion channel genes (KCNJ2, KCNJ4, HCN4, SCNA5, CACNA1C, CACNA1D, CACNA1G, KCNA5, KCNH2), TNNT2, and GAPDH was determined using single-cell real-time quantitative PCR (sc-RT-qPCR). For this approach, Ambiońs Single Cell-to-CT kit (Thermo Fisher Scientific, Waltham, MA, USA) in combination with the TaqMan gene expression assay technology (Thermo Fisher Scientific, Waltham, MA, USA) was used. The TaqMan gene expression assays are listed in Table 1. The experimental protocol was performed according to manufactureŕs instructions with the reagents contained in the kit. Briefly, the following steps were performed: After having collected single cells (see 2.4) in single nuclease-free 0.2 mL PCR-tubes (Eppendorf AG, Hamburg, Germany, already containing 9 µL Single Cell Lysis Solution and 1 µL Single Cell DNase), samples were incubated at rt for 5 min. Next, 1 µL Single Cell Stop Solution was added and samples were incubated at rt for 2 minutes and subsequently frozen (-20 °C) until further processing. 4.5 µL RT Master Mix was added to each sample and reverse transcription (25 °C for 10 min, 42 °C for 60 min, and 85 °C for 5 min) on a thermal cycler was performed to synthesize cDNA. Next, 11 µL PreAmp reaction mix containing pooled 0.2 x TaqMan Gene Expression Assays for all targets, was added to every sample and targeted preamplification was performed (95°C for 10 min, 14 cycles of 95°C for 15 s and 60°C for 4 min, and finally 99°C for 4 min). Pre-amplified samples were diluted 1:10 with 1 x TE Buffer (Thermo Fisher Scientific, Waltham, MA, USA). 10 µL 2 x TaqMan Gene Expression Master Mix, 4.0 µL pre-amplified and diluted product, 1.0 µL 20 X TaqMan Gene Expression Assay and 5.0 µL nuclease-free water were added per well of a 384-well plate. RT-qPCR (50 °C for 2 min, 95 °C 10 min, and 40 cycles of 95 °C for 5 s and 60 °C for 1 min) of every target was performed in single reactions as technical duplicates using the Roche Lightcycler 480 (Hoffmann-La Roche, Basel, Switzerland). A no-template-control (w/o cDNA) was included in all experiments as negative control. CT values (inversely proportional to the number of mRNA molecules on the log_2_ scale) were obtained using the 2^nd^-derivation method.

Detection of TNNT2 (structural cardiac gene) and GAPDH (potential house-keeping gene) was used to confirm the successful aspiration of a cell. Thus, samples not showing expression of TNNT2 or GAPDH were excluded from analysis. In case a CT value was given the tag “detector call uncertain“ (often the case for very high CT-values) or in case of atypical amplification curves, this CT was replaced by the value of the technical replicate (if available and valid) or excluded from analysis (if the replicate was invalid as well). In case the Lightcycler 480 did not detect a signal, reactions were indicated with “CT = 0”. The highest CT value reported by the instrument for our experiments was “CT = 35”, tagged with the note “Late Cp call (last five cycles) has higher uncertainty”. It was reported previously, that very high CT values may not be reliable and that the CT value corresponding to a single target cDNA molecule is usually 35-37 cycles for micro titer plates (Ståhlberg, Rusnakova, Forootan, Anderova, & Kubista, 2013). Therefore, we set all “CT = 0” values to “CT = 35” and defined a value of CT = 35 as “negative” result (no detectable expression). Accordingly, the percentage of “positive cells” was calculated for binary analysis. For further quantitative, statistical analysis, CT values of CT = 35 were excluded. A more accurate method would have been to determine a standard curve for every target gene and derive the slope (efficiency) and y intercept (detection limit). However, determining standard curves for single-cell experiments is challenging, as the calibrator sample should consist of the same complex biological matrix and be processed in exactly the same way as the experimental samples. As we did not determine standard curves/efficiencies, we performed downstream analysis using CT values and did not calculate numbers of transcripts. Moreover, we did not compare between different ion channels, as we cannot assure that amplification efficiencies were comparable between different targets. Consequently, we calculated the mean CT and standard deviation (SD) for the positive cells as surrogate for the expression level. Not surprisingly, for cells of different origins, we recognized that primary cardiomyocytes have overall lower CT values (higher expression levels) for every target gene investigated except HCN4 and CACNA1G than the three hiPSC-CM groups (see Table 3). The reason may be that primary cardiomyocytes are larger than hiPSC-CMs. However, individual normalization of single-cell data is generally not recommended by the manufacturer (*FAQ Single Cell-to-CT qRT-PCR Kit*, 2020) and in literature (Ståhlberg et al., 2013). Nevertheless, in order to compare expression between the five cell groups despite of their different expression levels, we normalized each individual CT value for a specific ion channel with respect to the mean CT value of GAPDH and TNNT2 of the whole cell group. We calculated ΔCT values for every cell and target gene by subtracting the averaged CT of TNNT2 and GAPDH for the whole cell group from the respective ion channel CT for each individual cell:

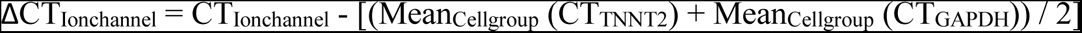

**Table 3:** Results of single-cell RT-qPCR. Besides the percentage of positive cells (% pos. Cells) and the number of investigated cells (n), also CT values are listed. Primary ventricular and atrial cardiomyocytes showed an overall higher expression level (lower CT values for every target gene investigated except HCN4 and CACNA1G) than the three hiPSC-CM groups. In order to compare expression between the five cell groups, despite their different sizes, we normalized each individual CT value for a specific ion channel with respect to the mean CT value of GAPDH and TNNT2 of the respective cell group. The resulting normalized CT values (=ΔCT) are listed in this table as well. Standard deviations (SD) are identical for CT and ΔCT values because the difference between the two values is a constant. For details of analysis of single-cell RT-qPCR data, see section 2.5 .

|  |  | HCN4 | CACNA1G | CACNA1D | KCNA5 | KCNJ4 | SCN5A | KCNJ2 | CACNA1C | KCNH2 | TNNT2 | GAPDH |
| --- | --- | --- | --- | --- | --- | --- | --- | --- | --- | --- | --- | --- |
| **iCell**<br>**CMs** | % pos. cells | 100 | 40 | 29 | 29 | 65 | 100 | 31 | 98 | 98 | 100 | 98 |
| | $\phi$ CT | 27.94 | 29.88 | 31.56 | 32.05 | 30.99 | 27.44 | 31.77 | 27.43 | 27.34 | 20.38 | 24.06 |
| | $\phi$ $\Delta CT$ | 5.59 | 7.53 | 9.22 | 9.70 | 8.64 | 5.10 | 9.42 | 5.08 | 4.99 | | |
| | ± SD | 1.49 | 1.89 | 1.56 | 1.46 | 1.29 | 1.61 | 1.26 | 1.81 | 1.47 | 1.02 | 1.17 |
|  | n | 55 | 55 | 55 | 55 | 55 | 55 | 55 | 55 | 55 | 55 | 55 |
| **Ventr.**<br>**PCs** | % pos. cells | 93 | 24 | 24 | 45 | 62 | 100 | 93 | 100 | 97 | 100 | 100 |
| | $\phi$ CT | 28.78 | 30.95 | 30.66 | 31.87 | 31.61 | 26.27 | 30.63 | 27.03 | 25.91 | 20.59 | 23.19 |
| | $\phi$ $\Delta CT$ | 6.89 | 9.06 | 8.77 | 9.99 | 9.72 | 4.39 | 8.74 | 5.14 | 4.02 | | |
| | ± SD | 1.19 | 1.20 | 1.45 | 0.85 | 1.38 | 1.62 | 1.98 | 1.60 | 1.49 | 1.11 | 1.20 |
|  | n | 29 | 29 | 29 | 29 | 29 | 29 | 29 | 29 | 29 | 29 | 29 |
| **Atr.**<br>**PCs** | % pos. cells | 100 | 63 | 67 | 96 | 42 | 100 | 58 | 92 | 100 | 100 | 100 |
| | $\phi$ CT | 26.52 | 29.43 | 30.34 | 28.72 | 31.82 | 26.80 | 30.15 | 27.27 | 26.23 | 19.67 | 23.44 |
| | mean $\Delta CT$ | 4.97 | 7.87 | 8.79 | 7.17 | 10.27 | 5.25 | 8.59 | 5.71 | 4.68 | | |
| | ± SD | 2.19 | 2.34 | 2.18 | 2.72 | 1.16 | 1.98 | 2.07 | 2.33 | 2.31 | 1.81 | 2.27 |
|  | n | 24 | 24 | 24 | 24 | 24 | 24 | 24 | 24 | 24 | 24 | 24 |
| **Prim.**<br>**ventr.**<br>**CMs** | % pos. cells | 94 | 9 | 14 | 88 | 91 | 100 | 97 | 100 | 100 | 100 | 100 |
| | $\phi$ CT | 28.58 | 29.84 | 29.48 | 29.01 | 28.95 | 24.05 | 27.18 | 25.68 | 24.48 | 16.69 | 20.91 |
| | $\phi$ $\Delta CT$ | 9.78 | 11.04 | 10.68 | 10.21 | 10.16 | 5.25 | 8.38 | 6.88 | 5.68 | | |
| | ± SD | 1.73 | 1.60 | 2.19 | 1.85 | 1.80 | 1.93 | 1.73 | 2.25 | 1.80 | 1.41 | 1.32 |
|  | n | 66 | 66 | 66 | 66 | 66 | 66 | 66 | 66 | 66 | 66 | 66 |
| **Prim.**<br>**atr.**<br>**CMs** | % pos. cells | 95 | 52.5 | 65 | 100 | 95 | 100 | 80 | 97.5 | 100 | 100 | 100 |
| | $\phi$ CT | 28.11 | 30.35 | 30.43 | 27.11 | 29.72 | 24.85 | 29.98 | 25.95 | 25.75 | 17.93 | 21.25 |
| | $\phi$ $\Delta CT$ | 8.52 | 10.76 | 10.84 | 7.52 | 10.13 | 5.25 | 10.39 | 6.35 | 6.16 | | |
| | ± SD | 2.02 | 1.39 | 1.84 | 1.85 | 1.73 | 1.44 | 1.93 | 1.63 | 1.67 | 1.37 | 1.32 |
|  | n | 40 | 40 | 40 | 40 | 40 | 40 | 40 | 40 | 40 | 40 | 40 |

The five cell groups were compared regarding the percentage of positive cells and the mean normalized expression level in these positive cells (ΔCT).

### 2.6 Statistics

SAS software (SAS institute, Cary, NC, USA) was used for statistical analysis. For every electrophysiological and expression parameter, mean and SD (per cell group) were calculated. GraphPad Prism (GraphPad software, San Diego, CA, USA) was used to create bar graphs. Spotfire (TIBCO, Somerville, MA, USA) software was used to create histograms.

Electrophysiological parameters and ΔCT values (see section 2.5) were compared between cell groups. Each parameter was analysed separately, using a linear model for repeated measurements (SAS proc mixed). “Cell group” was considered as a fixed effect. The covariance structure was modelled by a compound symmetry (CS) structure and the covariance parameters were estimated using residual (restricted) maximum likelihood (REML). The pairwise comparisons were made using Student’s t-test. The p-values were not adjusted for multiple comparisons.

Electrophysiological parameters and ΔCT values were compared between beating and non-beating cells for every cell group. Each parameter was analyzed separately. Beat rate was considered as a fixed effect in a linear model for repeated measurements. The covariance structure was modelled by a compound symmetry (CS) structure possibly different for the two groups. The covariance parameters were estimated using a residual (restricted) maximum likelihood procedure (REML).

To assess the relation between the single-cell parameters, each parameter was correlated with every other parameter (in total 18 parameters, see Table 6) and Pearsońs coefficient was used for quantification (positive coefficient indicates direct correlation, negative coefficient indicates inverse relationship, absolute value of the coefficient indicates strength of correlation).

**Table 6:** Parameters used in correlation analysis.

|  |  |
| --- | --- |
| Co-variables: | Capacitance [pF], CT of TNNT2, CT of GAPDH |
| Experimental parameters: | Spontaneous beat rate [bpm] (beating and non-beating cells), inter-beat interval [ms] (only beating cells), fast inward current amplitude [nA], fast inward current density [nA/pF], upstroke velocity [V/ms], APD90 [ms], CT of KCNJ4, CT of HCN4, CT of SCN5A, CT of CACNA1C, CT of KCNH2, CT of CACNA1D, CT of CACNA1G, CT of KCNA5, CT of KCNJ2 |

## 3 Results

### 3.1 Electrophysiological parameters: Variation within and differences between cell groups

A descriptive summary of the electrophysiological parameters is displayed in Figure 2 and listed in Table 2. For representative traces illustrating the range of action potentials, see Figure 1. The electrical capacitance, a surrogate parameter for plasma membrane surface area and therefore an indicator for cell size, was comparable between the three iPSC-CM groups, whereas human primary ventricular cardiomyocytes had a significantly higher capacitance. In line with our visual observation that single cells of the same type differed considerably in size, the variability in capacitance was high for all cell types, reflecting the range in the population. The action potential duration, fast inward current amplitude and fast inward current density were comparable between iCell cardiomyocytes and ventricular Pluricytes, without statistically significant differences. The values of these three parameters were generally smaller for atrial Pluricytes and primary human ventricular cardiomyocytes: Statistically significant differences can be observed between atrial Pluricytes and iCell cardiomyocytes/ventricular Pluricytes and primary ventricular cardiomyocytes and iCell cardiomyocytes/ventricular Pluricytes.

**Figure 1:**
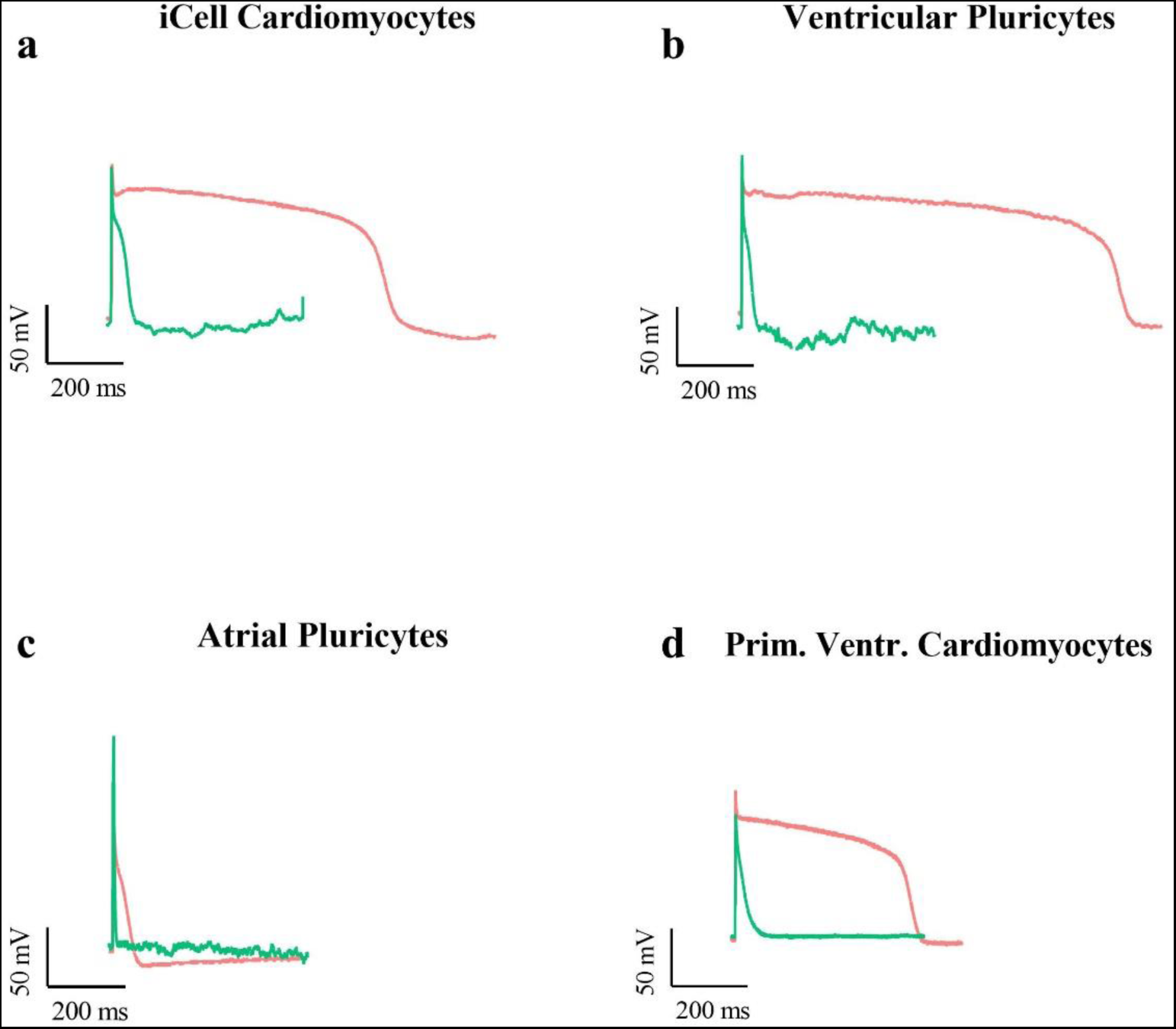
Representative action potential traces. To display the spectrum of action potential durations, a cell with a long and a cell with a short action potential for each cell group (**(a)** iCell cardiomyocytes, **(b)** ventricular and **(c)** atrial Pluricytes, **(d)** primary adult human ventricular cardiomyocytes) was selected. A short action potential is shown in green and a long action potential is shown in red. The numbers of total recordings are listed in Table 4. Note that electrophysiological recordings were not available from primary atrial cardiomyocytes.

**Figure 2:**
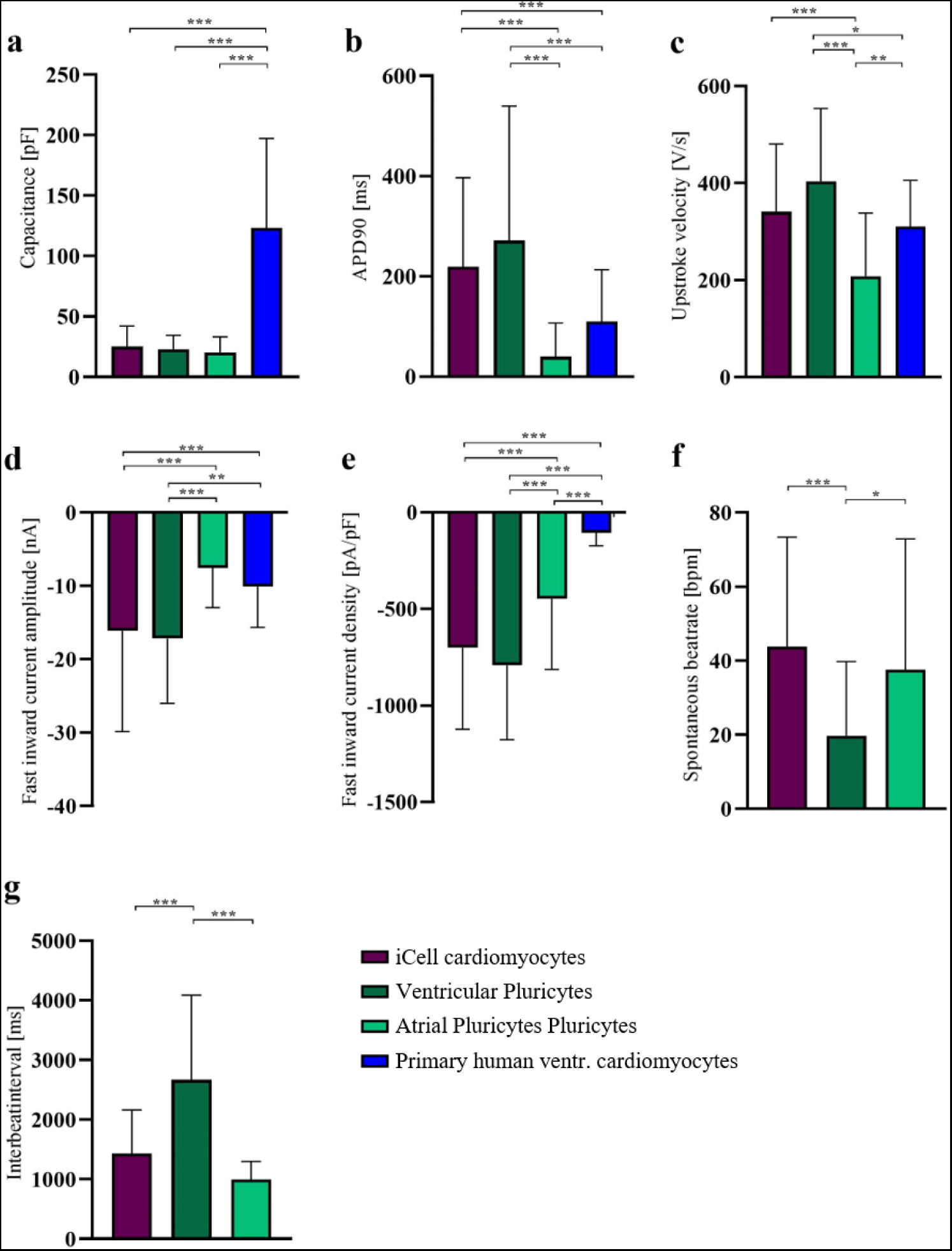
Comparison of electrophysiological parameters between cell groups. Data are shown as mean ± SD. Students t-test was used to compare groups. * indicates p<0.05; ** indicates p<0.01, and *** indicates p<0.001. The numbers of total recordings used to calculate means are listed in Table 4. Note that electrophysiological recordings were not available from primary atrial cardiomyocytes.

**Table 2:** Results of electrophysiological measurements. . Note that electrophysiological recordings were not available from primary atrial cardiomyocytes. Data are presented as means ± standard deviation (SD) and sample size (n).

|  |  | Capacitance<br>[pF] | APD90<br>[ms] | Spontaneous<br>beat rate<br>[bpm] | Inter-beat-<br>interval<br>[ms] | Upstroke<br>velocity<br>[V/s] | Fast inward<br>current<br>amplitude<br>[nA] | Fast inward<br>current<br>density<br>[pA/pF] |
| --- | --- | --- | --- | --- | --- | --- | --- | --- |
| iCell<br>CMs | mean | 25.15 | 219.00 | 43.75 | 1429.5 | 340.64 | -16.12 | -700.48 |
| | ± SD | 16.94 | 177.64 | 29.60 | 729.6 | 140.00 | 13.74 | 423.26 |
|  | n | 72 | 72 | 72 | 60 | 70 | 71 | 71 |
| Ventr.<br>PCs | mean | 23.00 | 271.36 | 19.75 | 2665.7 | 403.75 | -17.17 | -791.36 |
| | ± SD | 11.44 | 268.50 | 20.04 | 1418.3 | 150.00 | 8.85 | 385.25 |
|  | n | 32 | 32 | 32 | 21 | 30 | 29 | 29 |
| Atr.<br>PCs | mean | 20.36 | 39.81 | 37.58 | 994.4 | 208.43 | -7.55 | -447.24 |
| | ± SD | 12.91 | 66.89 | 35.30 | 305.1 | 130.00 | 5.41 | 366.18 |
|  | n | 33 | 29 | 33 | 19 | 27 | 27 | 27 |
| Prim.<br>ventr.<br>CMs | mean | 123.16 | 110.35 |  |  | 310.20 | -10.11 | -105.32 |
| | ± SD | 74.02 | 102.88 | | | 95.56 | 5.55 | 67.70 |
|  | n | 46 | 52 |  |  | 34 | 46 | 45 |

**Table 4:** Rank order of expression results. Cell groups are sorted starting with highest mean (ø) ΔCT (lowest expression) and smallest percentage of positive cells. Meaning of the symbols used for comparison are as described: Comparability: “≈” (no significant difference in ø ΔCT, difference in percentage of positive cells ≤ 15%), “>”: p<0.05, difference in percentage of positive cells >15%; “>>”: p<0.01; and “>>>”: p<0.001). Symbols are only indicating the degree of difference between one cell group and the following one.

|  |  |  |
| --- | --- | --- |
| HCN4 | % pos. Cells<br>$\phi$ ΔCT | ventr. PCs ≈ prim. ventr. CMs ≈ prim. atr. CMs ≈ atr. PCs ≈ iCells<br>prim. ventr. CMs $>>>$ prim. atr. CMs $>>>$ ventr. PCs $>>$ iCells ≈ atrial PCs |
| CACNA1G | % pos. Cells<br>$\phi$ ΔCT | prim. ventr. CMs ≈ ventr. PCs $<$ iCells ≈ prim. atr. CMs ≈ atr. PCs<br>prim. ventr. CMs ≈ prim. atr. CMs $>$ ventr. PCs ≈ atr. PCs ≈ iCells |
| CACNA1D | % pos. Cells<br>$\phi$ ΔCT | prim. ventr. CMs ≈ ventr. PCs ≈ iCells $<$ prim. atr. CMs ≈ atr. PCs<br>prim. atr. CMs ≈ prim. ventr. CMs ≈ iCells ≈ atr. PCs ≈ ventr. PCs |
| KCNA5 | % pos. Cells<br>$\phi$ ΔCT | iCells $<$ ventr. PCs $<$ prim. ventr. CMs ≈ atr. PCs ≈ prim. atr. CMs<br>prim. ventr. CMs ≈ ventr. PCs ≈ iCells $>>>$ prim. atr. CMs ≈ atr. PCs |
| KCNJ4 | % pos. Cells<br>$\phi$ ΔCT | Atr. PCs $<$ ventr. PCs ≈ iCells $<$ prim. ventr. CMs ≈ prim. atr. CMs<br>atr. PCs ≈ prim. ventr. CMs ≈ prim. atr. CMs ≈ ventr. PCs $>$ iCells |
| SCNA5 | % pos. Cells<br>$\phi$ ΔCT | prim. ventr. CMs ≈ ventr. PCs ≈ iCells ≈ prim. atr. CMs ≈ atr. PCs<br>prim. ventr. CMs ≈ prim. atr. CMs ≈ atr. PCs ≈ iCells ≈ ventr. PCs |
| KCNJ2 | % pos. Cells<br>$\phi$ ΔCT | iCells $<$ atr. PCs $<$ prim. atr. CMs ≈ ventr. PCs ≈ prim. ventr. CMs<br>prim. atr. CMs ≈ iCells ≈ ventr. PCs ≈ atr. PCs ≈ prim. ventr. CMs |
| CACNA1C | % pos. Cells<br>$\phi$ ΔCT | atr. PCs ≈ prim. atr. CMs ≈ iCells ≈ ventr. PCs ≈ prim. ventr. CMs<br>prim. ventr. CMs ≈ prim. atr. CMs ≈ atr. PCs ≈ ventr. PCs ≈ iCells |
| KCNH2 | % pos. Cells<br>$\phi$ ΔCT | ventr. PCs ≈ iCells ≈ prim. ventr. CMs ≈ prim. atr. CMs ≈ atr. PCs<br>prim. atr. CMs ≈ prim. Ventr. CMs $>$ iCells ≈ atr. PCs ≈ ventr. PCs |

When comparing APD90, fast inward current amplitude and density between atrial Pluricytes and primary ventricular cardiomyocytes, a significant difference was noted for the fast inward current density, with atrial Pluricytes having a higher mean density. The velocity of action potential upstroke was significantly slower in atrial Pluricytes compared to every other cell group. Moreover, upstroke velocity of primary ventricular cardiomyocytes was significantly slower than in ventricular Pluricytes. Differences of the spontaneous beat rate were only investigated in hiPSC-CMs, as primary ventricular cardiomyocytes generally do not beat without external stimulation. For a high percentage of single hiPSC-CMs, spontaneous beating was observed (iCell cardiomyocytes: 83 %, ventricular Pluricytes: 66 %, atrial Pluricytes: 58%). With roughly 40 bpm, iCell cardiomyocytes and atrial Pluricytes had a comparable intrinsic frequency whereas ventricular Pluricytes were significantly slower (roughly 20 bpm). Although results for beating behavior change slightly when only beating cells are used for calculations, ventricular Pluricytes consistently show a significantly higher inter-beat interval (30.10 ± 17.19 bpm / 2665.7 ± 1418.3 ms (n=21)) compared to the other two cell groups (iCell cardiomyocytes: 52.50 ± 24.23 bpm / 1429.5 ± 140 ms (n=60), atrial Pluricytes: 65.26 ± 17.55 bpm / 994.4 ± 305.1 ms (n=19)). All parameters showed high variation.

### 3.2 Comparison of expression between cell types

The aim of this study was to investigate both, electrophysiological parameters and expression of cardiac ion channels, in the same cell. Therefore, we performed single-cell RT-qPCR of nine cardiac ion channel subunits (HCN4, CACNA1G, CACNA1D, KCNA5, KCNJ4, SCN5A, KCNJ2, CACNA1D and KCNH2), the cardiac related gene TNNT2 and the potential housekeeping gene GAPDH using hiPSC-CMs (iCell cardiomyocytes, ventricular Pluricytes and atrial Pluricytes) and primary human ventricular and atrial cells. For a comprehensive summary, binary (positive/negative signal in single-cell RT-qPCR) expression profiles of every single cell investigated are displayed in Figure 3 (for more information on data analysis of single-cell RT-qPCR data see section 2.5). The quantitative results of the single-cell RT-qPCR are displayed in Figure 4 and listed in Table 3. Statistically significant differences of the pairwise comparisons of ΔCT values are shown in Figure 4. To compare expression between the cell groups despite their different expression levels, ΔCT values were calculated (CT values (of the positive cells) normalized to the mean expression of TNNT2 and GAPDH of the respective cell group). For details, see section 2.5.

**Figure 3:**
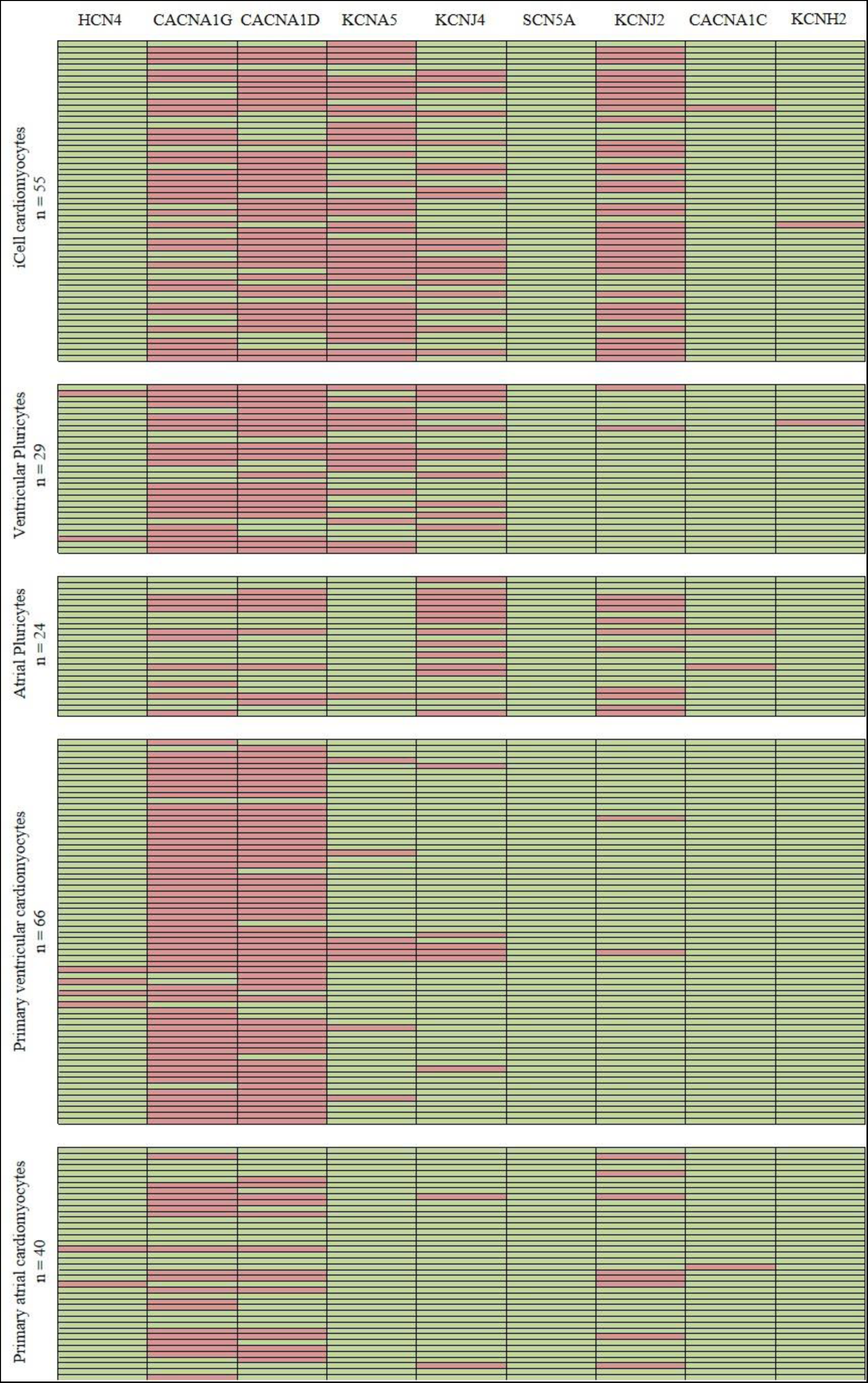
Binary expression analysis displays co-expression of cardiac ion channels in single CMs. Each row represents one cell. A green/red box indicates that the cell was positive/negative for the respective ion channel (for more information on data analysis of single-cell RT-qPCR data see section 2.5). Whenever APD was recorded, rows are sorted with increasing APD.

**Figure 4:**
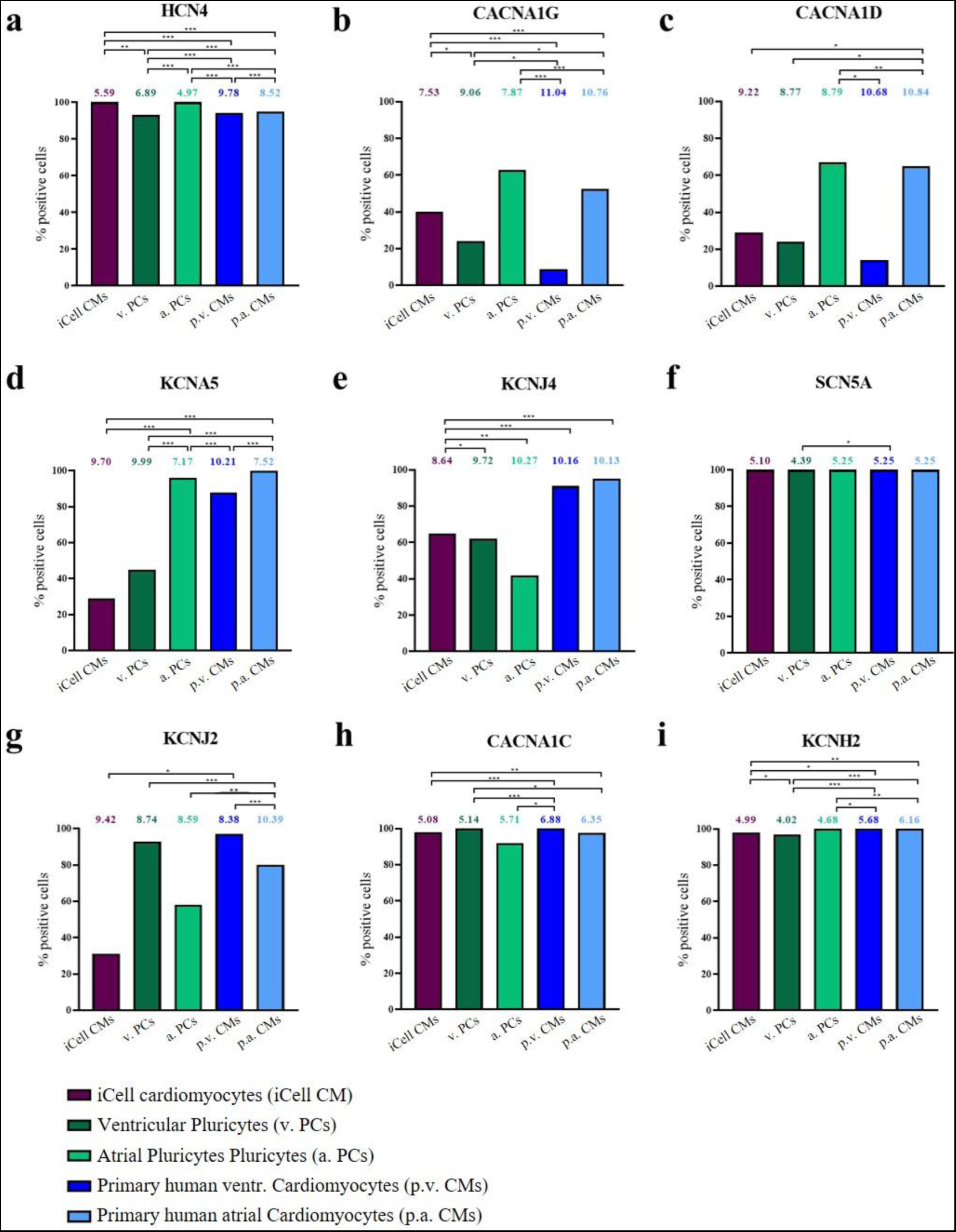
Comparison of cell groups regarding cardiac ion channel expression on single-cell level. Columns represent percentages of positive cells, values indicated above are mean ΔCT values (CT values normalized to the mean expression of TNNT2 and GAPDH of the respective cell group) of the positive cells and significant differences of ΔCT values between groups are indicated with p<0.05: *; p<0.01: ** and p<0.001: ***. The total numbers used to calculate mean ΔCT values are listed in Table 5.

**Table 5:** Comparison of spontaneously beating and non-beating hiPSC-CMs. In case the difference in percentage of positive cells (% pos. cells) between beating and non-beating hiPSC-CMs is >15 % boxes are highlighted with a yellow shading. In case of a significant difference between beating and non-beating hiPSC-CMs (p<0,05: *; p<0,01: ** and p<0.001: ***), values (mean (ø) CT, electrophysiological parameters) are written in blue. As comparisons were done within each cell group, CT values (and not ΔCT values) were used. However, the results for ΔCT values would be the same as for CT values, because the difference between the two values is a constant. The covariance parameters were estimated using residual (restricted) maximum likelihood procedure (REML).

|  |  | iCell Cardiomyocytes |  | Ventricular Pluricytes |  | Atrial Pluricytes |  |
| --- | --- | --- | --- | --- | --- | --- | --- |
| Spontaneous beat rate (n cells) |  | =0 (8) | >0 (47) | =0 (11) | >0 (18) | =0 (12) | >0 (12) |
| HCN4 | % pos. cells | 100 | 100 | 91 | 94 | 100 | 100 |
| | $\phi$ CT ± SD | 27.8 ± 0.6 | 28.0 ± 0.2 | 28.6 ± 0.4 | 28.9 ± 0.3 | 27.3 ± 0.7 | 25.8 ± 0.5 |
| CACNA1G | % pos. cells | 25 | 43 | 18 | 28 | 50 | 75 |
| | $\phi$ CT ± SD | 30.1 ± 0.1 | 29.9 ± 0.4 | 32.1 ± 0.5 | 30.5 ± 0.5 | 30.4 ± 1.0 | 28.8 ± 0.7 |
| CACNA1D | % pos. cells | 13 | 32 | 27 | 22 | 50 | 83 |
| | $\phi$ CT ± SD | 29.5 ± 0.03 | 31.7 ± 0.4 *** | 31.0 ± 0.9 | 30.4 ± 0.8 | 31.3 ± 0.8 | 29.8 ± 0.7 |
| KCNA5 | % pos. cells | 38 | 28 | 64 | 33 | 92 | 100 |
| | $\phi$ CT ± SD | 32.6 ± 0.4 | 31.9 ± 0.4 | 31.7 ± 0.3 | 32.1 ± 0.4 | 29.5 ± 0.9 | 28.0 ± 0.7 |
| KCNJ4 | % pos. cells | 75 | 64 | 64 | 61 | 42 | 42 |
| | $\phi$ CT ± SD | 30.8 ± 0.6 | 31.0 ± 0.2 | 31.9 ± 0.6 | 31.4 ± 0.4 | 32.2 ± 0.5 | 31.5 ± 0.5 |
| SCN5A | % pos. cells | 100 | 100 | 100 | 100 | 100 | 100 |
| | $\phi$ CT ± SD | 27.9 ± 0.4 | 27.4 ± 0.2 | 26.5 ± 0.5 | 26.1 ± 0.4 | 27.4 ± 0.5 | 26.2 ± 0.6 |
| KCNJ2 | % pos. cells | 25 | 32 | 100 | 89 | 50 | 67 |
| | $\phi$ CT ± SD | 31.6 ± 1.2 | 31.8 ± 0.3 | 30.4 ± 0.5 | 30.8 ± 0.5 | 31.0 ± 0.8 | 29.5 ± 0.7 |
| CACNA1C | % pos. cells | 100 | 98 | 100 | 100 | 83 | 100 |
| | $\phi$ CT ± SD | 27.7 ± 0.9 | 27.4 ± 0.3 | 26.5 ± 0.4 | 27.4 ± 0.4 | 28.3 ± 0.9 | 26.4 ± 0.5 |
| KCNH2 | % pos. cells | 100 | 98 | 91 | 100 | 100 | 100 |
| | $\phi$ CT ± SD | 27.9 ± 0.7 | 27.2 ± 0.2 | 25.4 ± 0.2 | 26.2 ± 0.4 | 27.0 ± 0.7 | 25.5 ± 0.6 |
| $\phi$ Capacitance [pF] ± SD | | 14.68 ± 1.80 | 27.25 ± 2.28 *** | 21.35 ± 3.14 | 23.86 ± 2.64 | 21.24 ± 4.39 | 19.71 ± 2.30 |
| $\phi$ APD90 [ms] ± SD | | 132 ± 42 | 236 ± 23 * | 222 ± 72 | 304 ± 34 | 54 ± 29 | 31 ± 10 |
| $\phi$ Fast inward current amplitude [nA] ± SD | | -10.36 ± 2.33 | -17.30 ± 1.87 * | -12.05 ± 1.12 | -19.86 ± 2.22 ** | -6.82 ± 1.56 | -7.98 ± 1.40 |
| $\phi$ Fast inward current density [pA/pF] ± SD | | -686.1 ± 125.8 | -703.4 ± 55.3 | -607.8 ± 72.2 | -888.0 ± 96.3 * | -450.9 ± 111.8 | -445.1 ± 93.2 |
| $\phi$ Upstroke velocity [V/ms] ± SD | | 0.3119 ± 0.03583 | 0.3461 ± 0.01924 | 0.2952 ± 0.0451 | 0.4662 ± 0.0279 ** | 0.1789 ± 0.0425 | 0.2260 ± 0.0321 |

#### 3.2.1 Transcripts of SCN5A, CACNA1C, KCNH2 and HCN4 are found in a high percentage of cardiomyocytes of every cell group

Transcripts of four ion channels (SCN5A, CACNA1C, KCNH2 and HCN4) were found in almost every cell (92-100%) of every cell group. The mean expression levels (ΔCT) for SCN5A were comparable between the groups. Only between ventricular Pluricytes and ventricular primary cardiomyocytes a significant difference could be observed, indicating that the mean expression in ventricular Pluricytes was higher (lower ΔCT value) than in ventricular primary cardiomyocytes (see Figure 4, Table 3). For CACNA1C, no significant difference between the two primary cardiomyocyte groups and between the three hiPSC-CM groups could be observed. However, pairwise comparisons between primary atrial or ventricular cardiomyocytes and iCell cardiomyocytes or ventricular Pluricytes or atrial Pluricytes, revealed significantly higher ΔCT values (lower expression) for the primary cells (except for atrial Pluricytes and primary atrial cardiomyocytes).

Also, for KCNH2, ΔCT values were comparable between primary atrial and ventricular cardiomyocytes. Pairwise testing of primary atrial or ventricular cardiomyocytes with iCell cardiomyocytes or ventricular Pluricytes or atrial Pluricytes, respectively, revealed significantly higher ΔCT values (lower expression) for the primary cells. Moreover, mean ΔCT was significantly higher (lower expression) for iCell cardiomyocytes compared to ventricular Pluricytes. Significant differences in mean ΔCT of HCN4 can be found in every pairwise comparison except for iCell cardiomyocytes and atrial Pluricytes. The highest mean ΔCT (lowest expression) can be found in primary ventricular cardiomyocytes, followed by primary atrial cardiomyocytes, ventricular Pluricytes, iCell cardiomyocytes and atrial Pluricytes (lowest mean ΔCT/highest expression).

#### 3.2.2 Higher expression (ΔCT) of CACNA1G, CACNA1D and KCNA5 in atrial compared to ventricular cells

For CACNA1G and CACNA1D, the same trends regarding the percentage of positive cells was observed: A majority of atrial cardiomyocytes (Pluricytes and primary cardiomyocytes, 53-67%) was positive, whereas the percentage in their ventricular counterparts was much smaller (9-24%). The percentage of positive cells for iCell cardiomyocytes was between the percentage of atrial and ventricular cells. Pairwise comparisons of mean ΔCTs for CACNA1G revealed significant differences between primary atrial or ventricular cardiomyocytes with each of the three hiPSC-CM groups.

In addition, ventricular Pluricytes showed a significantly higher mean ΔCT than iCell cardiomyocytes. For CACNA1D, significant differences were seen: The mean ΔCT of primary atrial cardiomyocytes was significantly higher (lowest expression) than the mean ΔCT of the three hiPSC-CM groups. Moreover, there is a significant difference between atrial Pluricytes (lower mean ΔCT) and primary ventricular cardiomyocytes (higher mean ΔCT). For KCNA5, 96% of the atrial Pluricytes and 100% of the primary atrial cardiomyocytes were positive with comparable mean ΔCTs. Both these cell groups show, compared to every other cell group, the significantly lowest mean ΔCT value (highest expression). Interestingly, with 88% also a high percentage of primary ventricular cardiomyocytes was positive for KCNA5.

#### 3.2.3 Expression of KCNJ2 and KCNJ4

The percentage of positive cells for KCNJ2 showed the same pattern for Pluricytes and primary cells: A higher percentage of ventricular Pluricytes and ventricular primary cardiomyocytes (93% and 97%, respectively) was positive for KCNJ2 than of their atrial counterpart (atrial Pluricytes: 31%, primary atrial cardiomyocytes: 80%). No significant difference was observed between the three hiPSC-CM groups for the mean ΔCT of KCNJ2. The mean ΔCT value of the primary atrial cardiomyocytes is significantly higher (lower expression) than the mean ΔCT of primary ventricular cardiomyocytes and ventricular Pluricytes. In addition, significant differences can be observed between primary atrial cardiomyocytes (higher mean ΔCT of KCNJ2) and atrial Pluricytes and iCell cardiomyocytes (higher mean ΔCT) and primary ventricular cardiomyocytes. The percentage of positive cells for KCNJ4 is higher in primary cardiomyocytes (ventricular: 91%, atrial: 95%) than in hiPSC-CMs (42-65%). The mean ΔCT values are comparable between the cell groups except for iCell cardiomyocytes: iCell cardiomyocytes have a significantly lower ΔCT value than every other cell group.

#### 3.2.4 Summary of single-cell expression results

To provide an intuitive overview of single-cell expression results, rank orders of mean ΔCT values and percentages of positive cells for every ion channel are listed in Table 4.

### 3.3 Comparison of spontaneously beating and non-beating hiPSC-CMs

A high percentage of single hiPSC-CMs showed spontaneous contractions (iCell cardiomyocytes: 83%, ventricular Pluricytes: 66%, atrial Pluricytes: 58%). However, there is also a subset of cells, which – when the spontaneous beating rate was to be counted immediately before patch clamping commenced – did not beat spontaneously. We hence asked whether beating and non-beating cells within one hiPSC-CM group (iCell cardiomyocytes, ventricular Pluricytes and atrial Pluricytes) differed with regard to ion channel expression (HCN4, CACNA1G, CACNA1D, KCNA5, KCNJ4, SCN5A, KCNJ2, CACNA1C and KCNH2) or electrophysiology (capacitance, APD90, fast inward current amplitude and fast inward current density). Therefore, we compared the percentage of positive cells for each ion channel and evaluated the differences in CT values and electrophysiology parameters between beating and non-beating cells within one cell group. As comparisons were done within each cell group, CT values (and not ΔCT values) were used for pairwise statistical testing. The results of this analysis are listed in Table 5. Statistical analysis in some cases needs to be taken with caution as *n* is small for some analyses (note “n” and “%” in Table 5).

No statistically significant differences in CT values can be observed between beating and non-beating cells within one cell group, except for CACNA1D: In iCell cardiomyocytes, non-beating cells showed a significantly smaller CT value (higher expression) than their beating counterpart. Percentage of positive cells is comparable between beating and non-beating cells of all three cell groups investigated for HCN4, KCNJ4, SCN5A and KCNH2. The percentage of CACNA1G- and CACNA1D-positive cells is higher in beating compared to non-beating iCell cardiomyocytes and atrial Pluricytes. Moreover, in atrial Pluricytes, the percentage of positive cells for KCNJ2 and CACNA1C is slightly higher in beating cells. In ventricular Pluricytes, the percentage of positive cells for KCNA5 is higher in non-beating cells. Regarding the electrophysiological parameters, beating iCell cardiomyocytes show a significantly higher capacitance and faster inward current amplitude and a longer APD90. In addition, ventricular Pluricytes show a higher fast inward current amplitude, density and upstroke velocity compared to non-beating cells.

### 3.4 Histograms of single-cell data

We constructed histograms to visualize the distribution of single-cell data (see Figure 5 and SuppFigure 1). Earlier publications claimed that 54% or 64% of iCell cardiomyocytes have ventricular-like action potentials, 22% or 18% have nodal-like action potentials, and 24% or 18% have atrial-like action potentials (according to Ma et al., 2011 or Yonemizu et al., 2019, respectively). If iCell cardiomyocytes consisted of nodal-atrial- and ventricular-like cardiomyocytes, distributions of subtype-specific parameters are supposed to be multiphasic, due to the assumed sub-populations. To test the hypothesis that histograms should reveal cardiac subpopulations, we first pooled our expression data (ΔCT) of atrial and ventricular primary human cardiomyocytes and asked whether the histograms would allow the discrimination of the primary subtypes (Figure 5a and b). Figure 5 shows histograms for ΔCT of KCNA5. It is known that KCNA5 (Potassium Voltage-Gated Channel Subfamily A Member 5, K_V_1.5, conducting I_Kur_) is predominantly expressed in primary atrial cardiomyocytes as compared to their ventricular counterpart (Kane & Terracciano, 2017). Figure 5a demonstrates that the two subpopulations can be recognized as long as data sets are separated (see Figure 5a: KCNA5: atrial primary cardiomyocytes are left-shifted compared to primary ventricular cardiomyocytes). However, when the two data sets are pooled (Figure 5b), differences between atrial and ventricular primary cardiomyocytes are too small to be resolved (for our number of investigated cells). Therefore, we do not anticipate that histograms of expression data from iCell cardiomyocytes are suitable to confirm three subpopulations.

**Figure 5:**
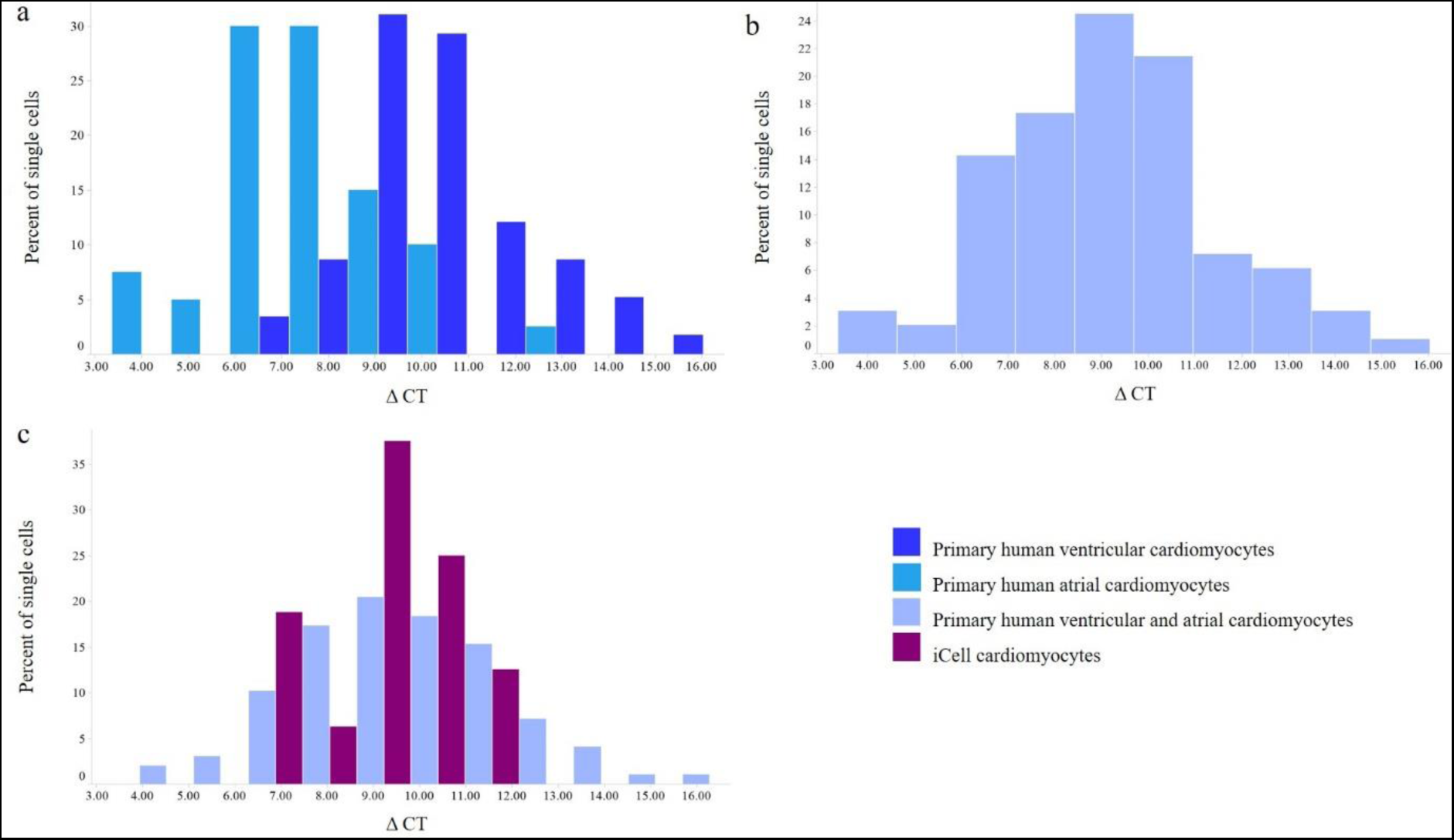
Histograms of ΔCT of KCNA5. Evaluation of histograms to confirm cardiac subpopulations. In (a) the separated data sets of primary atrial and ventricular cardiomyocytes clearly show the expected left-shifted distribution of atrial cardiomyocytes (higher expression of this atrial-associated gene). However, the histogram of the pooled data sets in (b) demonstrates that differences between atrial and ventricular primary cardiomyocytes are too small to be resolved (for our number of investigated cells). Thus, it is unlikely that histograms of expression data from iCell cardiomyocytes (c) are suitable to confirm three subpopulations. For total *n* see Table 5. The histograms of the other ion channels investigated can be found in SuppFigure 1.

For the electrophysiological parameters, data are only available for primary ventricular cardiomyocytes and we can therefore not evaluate whether shapes of histograms of electrophysiological parameters reflect cardiac subpopulations. However, the histograms of electrophysiological data from iCell cardiomyocytes do not indicate subpopulations, but show a bigger range of distribution (except for capacitance) compared to primary ventricular cardiomyocytes (see Figure 6).

**Figure 6:**
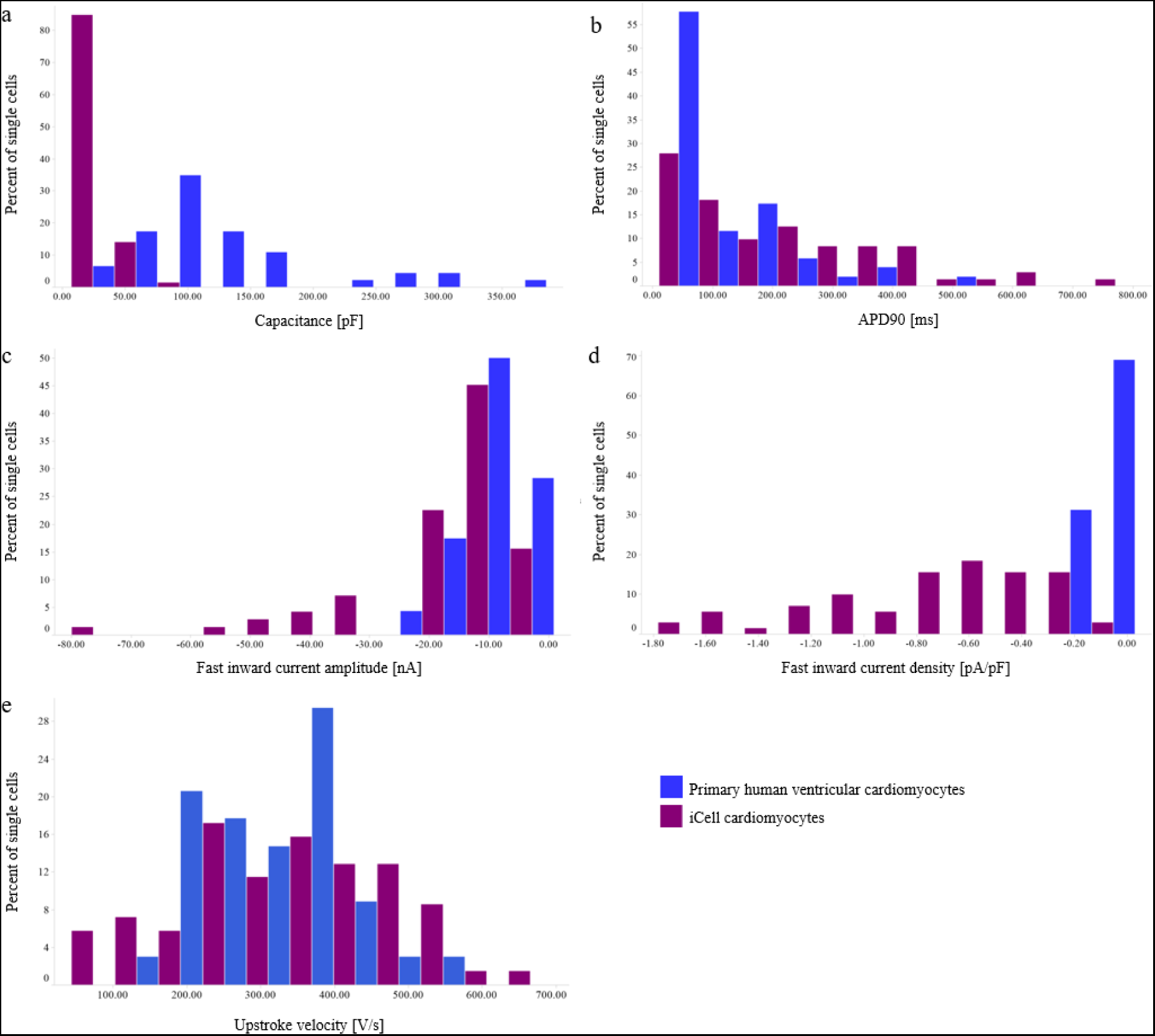
Histograms of electrophysiological parameters for iCell cardiomyocytes and primary human ventricular cardiomyocytes. As data of electrophysiological parameters are only available for primary ventricular cardiomyocytes (and not for primary atrial cardiomyocytes), we cannot evaluate whether shapes of histograms of electrophysiological parameters mirror cardiac subpopulations. However, the histograms of electrophysiological data from iCell cardiomyoctes do not indicate subpopulations. The total range of values is bigger than for primary ventricular cardiomyocytes (except for capacitance). For total *n* see Table 4.

### 3.5 Correlations on the single-cell level

To investigate the relation between all parameters determined on single-cell level, we correlated each parameter with every other parameter (in total 18 parameters, see Table 6) for each cell group investigated.

The results are shown in Supp. Table 2 - Supp. Table 6. Statistically significant (p < 0.05) correlations are highlighted with a light orange shading. Selected correlations of specific interest are presented in Figure 6. To provide a first insight into the complex data set, the following text lists all significant correlations and the expected but not significant correlations. The results are structured into correlations with capacitance, correlations of expression parameters with each other, correlations of expression and electrophysiology parameters and correlations of electrophysiological parameters with each other. For interpretations and physiological context, see section 4. Positive correlations are presented in **bold letters** and inverse correlations are indicated *italicized*.

#### 3.5.1 Several correlations with capacitance

Based on the results, especially for the three hiPSC-CM groups, several expression parameters *inversely correlate* significantly with the capacitance (iCells: *CT of TNNT2, GAPDH, SCN5A, KCNH2, CACNA1D and CACNA1G*. Ventricular Pluricytes: *CT of TNNT2, GAPDH, HCN4. SCN5A and KCNH2*. Atrial Pluricytes: *CT of GAPDH and KCNA5*). The higher the capacitance, the smaller the CT value (higher expression). For primary ventricular cardiomyocytes, capacitance only correlates significantly with *CT of KCNA5*. Regarding the hiPSC-CMs, some electrophysiology parameters also correlate with capacitance (iCell cardiomyocytes: **Beat rate**, *inter-beat interval*, *fast inward current amplitude*. Ventricular Pluricytes: *Fast inward current amplitude*, **fast inward current density**. Atrial Pluricytes: **Fast inward current density**, **APD90**). For primary ventricular cardiomyocytes, capacitance correlates with the **fast inward current density**.

#### 3.5.2 Expression parameters correlate with each other

Correlating expression parameters (CT values) against each other revealed strong and highly significant, **positive** correlations for many parameter pairs. This holds true for all five investigated cell groups – hiPSC-CMs as well as primary cardiomyocytes. Representative correlations are shown in SuppFigure 2, for details see SuppTable 2 - SuppTable 6. Besides these positive correlations, also few *inverse correlations* were observed (ventricular Pluricytes: *KCNA5 with KCNJ4, CACNA1D and CACNA1G*. Primary ventricular cardiomyocytes: *CACNA1D with HCN4 and CACNA1C*). For all inversely correlating pairs, *n* is low (n<10).

#### 3.5.3 Correlation of CT of SCN5A and fast inward current amplitude in hiPSC-CMs

A question of specific interest was whether expression and electrophysiological parameters correlated on a single-cell level, or, more general, whether the investigated transcriptomic phenotype can be related to the electrophysiological phenotype of individual cells. For all three hiPSC-CM groups, **CT of SCN5A** correlated positively and significantly with the **fast inward current amplitude** (see Figure 7). Further correlations between expression and electrophysiological parameters could be established (iCell cardiomyocytes: *Beat rate with KCNH2 and CACNA1G*, **inter-beat interval with CACNA1D**, fast **inward current amplitude with GAPDH**, fast **inward current density** and *APD90* with KCNJ2, **upstroke velocity with KCNJ4**. Ventricular Pluricytes: **Fast inward current amplitude with KCNH2 and CACNA1G**, *fast inward current density with GAPDH/KCNA5*. Atrial Pluricytes: *Beat rate with SCN5A*, **fast inward current amplitude with KCNJ4**). For primary ventricular cardiomyocytes, *CT of KCNA5 correlates with the fast inward current amplitude and density*. Some correlations (e.g. HCN4, KCNJ2, KCNJ4 with beat rate, KCNH2, CACNA1C with APD90), were expected due to a functional relationship but no correlation could be established.

**Figure 7:**
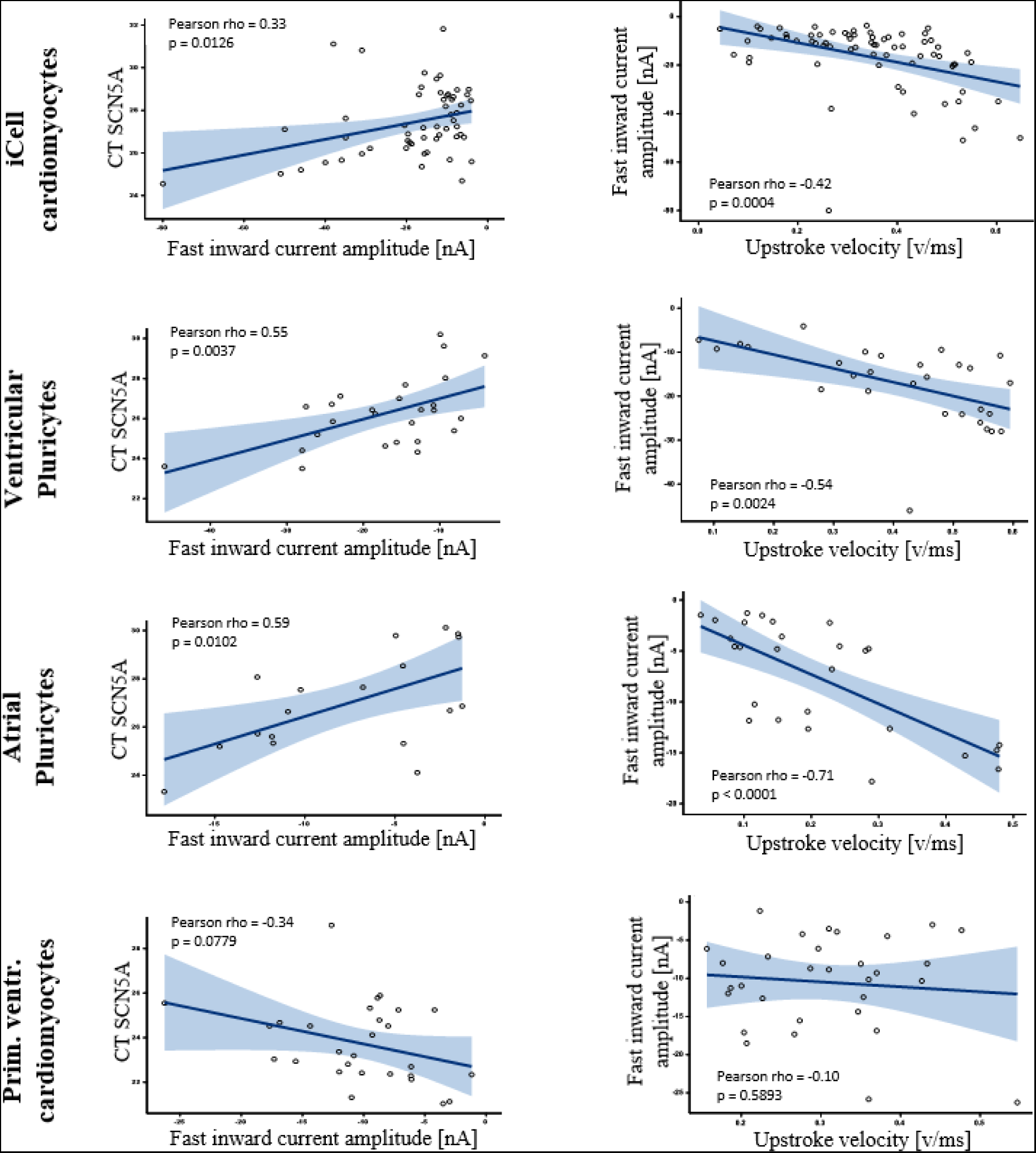
Selected single-cell correlations. For all three hiPSC-CM groups, correlations of CT SCN5A and fast inward current amplitude and fast inward current amplitude and upstroke velocity was observed. Blue shaded area indicates 95% confidence limits.

#### 3.5.4 Correlations between electrophysiological parameters

Besides expression-expression- and expression-electrophysiology-correlations, also electrophysiology-electrophysiology-correlations were found for hiPSC-CMs (but not for primary ventricular cardiomyocytes): For all three hiPSC-CM groups, an *inverse correlation* for the *fast inward current amplitude and density with the upstroke velocity* of the action potential was found (see Figure 7). Moreover, for iCell cardiomyocytes, *beat rate and fast inward current amplitude*, **inter-beat interval and APD90** were correlated. For ventricular Pluricytes, upstroke velocity correlated with **beat rate**.

## 4 Discussion

In the following section, the results for the different electrophysiological parameters together with ion channel expression results related to the respective parameter are discussed one after another. Moreover, we discuss the comparison of beating and non-beating hiPSC-CMs and whether histograms of single-cell parameters allow conclusions on possible subpopulations or whether single-cell phenotypes of hiPSC-CMs could be categorized into primary cardiac subtypes (nodal, atrial, and ventricular). Finally, we give implications for use.

### 4.1 Study design

iCell cardiomyocytes, ventricular Pluricytes, atrial Pluricytes, primary adult ventricular human cardiomyocytes and primary adult atrial human cardiomyocytes were investigated using a single-cell patch-clamp-RT-qPCR technique. As this method is technically challenging, it was not always possible to determine every parameter for every single cell. Therefore, *n* may vary between parameters. To enable the combination of electrophysiological measurements and subsequent RT-qPCR analysis, it is important to work quickly to minimize RNA loss (degradation, diffusion into the patch-pipette). Therefore, an in-depth pharmacological electrophysiological characterization was out of scope. However, we determined the capacitance, APD90 and upstroke velocity of the action potential, spontaneous beat rate, inter-beat interval, fast inward current amplitude, and fast inward current density of single cells. We showed in iCell cardiomyocytes that the fast inward current is mainly conducted by the cardiac sodium channel (NaV1.5, SCN5A, see SuppFigure 3a). For the subsequent expression analysis, the use of single-cell RNA-sequencing would have had the advantage of obtaining the whole transcriptome of single cells. However, in a previous study, we failed to detect ion channel transcripts using this method (Schmid et al., 2020). As other studies demonstrated that single-cell RT-qPCR is sensitive enough to detect ion channel expression (Burridge et al., 2014; Liang et al., 2013), we chose to employ this approach. With this method, we were able to detect ion channel transcripts but needed to limit (technical reasons: limited material in single cells) the transcripts of interest to nine important cardiac ion channel subunits (HCN4, CACNA1G, CACNA1D, KCNA5, KCNJ4, SCN5A, KCNJ2, CACNA1D and KCNH2). These ion channels were selected due to their anticipated relation to the determined electrophysiological parameters and/or due to their anticipated cardiac subtype associated expression (see introduction).

### 4.2 Reliability, robustness and limitations

Reliability and robustness of single-cell qPCR-data is indicated by highly similar CT values of technical duplicates (see SuppFigure 3d), demonstration of a high inter-plate reproducibility (see SuppFigure 3b) and reflection of cDNA dilutions in CT values (see SuppFigure 3d). Further technical details that strengthen the reliability of the single-cell RT-qPCR data are the application of sequence-specific taqman assays (which span introns whenever possible) and the reduction of risk for contamination by a one-tube-protocol without the need for mRNA-purification, DNAse treatment (to prevent gDNA impurification), dUTP/UDG application and a non-template-negative control. However, limitations that come along with this RT-qPCR approach are that we only assess expression of certain ion channel subunits on the transcriptomic level, which may not reflect functional ion currents and the complex electrophysiological phenotype of a cell (mRNA-protein-translation, other ion channel subunits and regulatory mechanisms, ion distributions, etc.). The patch-clamp rig on the one hand enabled single-cell electrophysiology measurements and harvesting of single cells, on the other hand it needs to be kept in mind that singularizing of cardiomyocytes and the selection of single cells may influence results (Du, Hellen, Kane, & Terracciano, 2015; Kane, Du, Hellen, & Terracciano, 2016). However, we observed a correlation between the mean single-cell RT-qPCR data and bulk-sequencing data of iCell cardiomyocytes (see SuppFigure 3c). In general, electrophysiological data are hard to compare between studies because of different protocols and experimental conditions. The interpretation of the single-cell correlations was challenging because of the complex data set (18 x 18 parameters), and the risk of indirect correlations.

The aim of our study was to present a transcriptional and electrophysiological snapshot of widely used cardiomyocyte models at an age when the cells are commonly used for cardiac safety assessment of novel drugs. Limitations that come along with this study design are that generated hiPSC-CM results are only valid for iCell and Pluricyte cardiomyocytes at the described age and cannot be transferred to hiPSC-CMs in general, as hiPSC-CMs seem to be highly variable depending on e.g. donor, culture conditions or age (Biendarra-Tiegs, Secreto, & Nelson, 2019). Thus, to provide the most comprehensive comparison for the iCell cardiomyocyte data, we primarily refer to the Ma et al., 2011 publication which systematically investigated various cardiac ion channel and currents. Although the period for utilization of hiPSC-CMs is relatively long (considering the technically challenging and time-consuming method), the number of days since thawing (for hiPSC-CMs) or days since singularizing (for hiPSC-CMs: singularizing from monolayer syncytium culture, for primary cardiomyocytes: Singularizing/isolation of single cells from heart tissue), did not turn out to be influential co-variables for the investigated parameters. This is in line with a correlation between bulk data (cells were lysed after two weeks in culture) and mean single-cell RT-qPCR data (see SuppFigure 3c).

### 4.3 Discussion of results

#### 4.3.1 **Capacitance**

When working with single cardiomyocytes, we visually observed an immense variation of cell sizes. Moreover, we observed - in accordance with previous publications - that hiPSC-CMs are small (Karakikes et al., 2015), i.e. smaller than adult primary human cardiomyocytes. In fact, the mean capacitance of iCell cardiomyocytes, ventricular Pluricytes and atrial Pluricytes was comparable (≈ 20-25 pF) and smaller than the mean capacitance of primary human ventricular cardiomyocytes (≈ 123 pF). This finding is in agreement with earlier reports (see collected data of human cardiomyocytes in (Polak & Fijorek, 2012) and data for hiPSC-CMs (Mann et al., 2019). Ventricular Pluricytes showed a slightly higher capacitance than atrial Pluricytes. This trend was also reported in other ventricular and atrial hiPSC-CMs (Cyganek et al., 2018; Pei et al., 2017; Zhang et al., 2011). The high standard deviation reported in our study for all cell types is in accordance with the observed morphological heterogeneity in cell size. Regarding single-cell correlations, many CT values of the investigated ion channel genes were found to inversely correlate with capacitance. This suggests that the single-cell expression depends on cell size. This trend is mainly seen in hiPSC-CMs. A plausible explanation could be that hiPSC-CMs (which are rather round and flat) have an active membrane surface area (and therefore capacitance) that correlates with cell volume compared to primary cardiomyocytes which exhibit a considerable membrane surface area enlargement due to the T-tubular system.

#### 4.3.2 Spontaneous beat rate, related ion channel expression (HCN4, KCNJ2, KCNJ4) and the comparison between spontaneously beating and non-beating hiPSC-CMs

The spontaneous firing of action potentials is a specific feature of pacemaker cardiomyocytes, like nodal cardiomyocytes from the sinoatrial (SA)/atrioventricular node. A critical difference between hiPSC-CMs and human primary atrial and ventricular cardiomyocytes is the spontaneous beating behavior of hiPSC-CMs. For a high percentage of single hiPSC-CMs, spontaneous beating was observed. For iCell cardiomyocytes, ventricular, and atrial Pluricytes we observed a mean spontaneous beat rate (beating and non-beating cells were included in calculation of mean beat rate) of ≈ 20-44 bpm, with iCell cardiomyocytes showing a significantly higher beat rate than ventricular Pluricytes. Our findings with single cells are comparable to those previously reported for hiPSC-CM monolayers: In multi-electrode array (MEA) experiments, iCell cardiomyocyte monolayers showed a spontaneous beat rate of 45 bpm and ventricular Pluricytes of 13 bpm (Rast et al., 2016). Moreover, atrial Pluricytes showed a significantly higher spontaneous beating frequency than ventricular Pluricytes, which is in line with findings from other atrial/ventricular hiPSC-CMs (Cyganek et al., 2018; Pei et al., 2017). Notably, values of beating frequency change slightly when the mean beat rate/inter-beat interval is calculated only for the beating cells. Taking this into consideration, ventricular Pluricytes still show the slowest beat rate/highest inter-beat interval. One likely mechanism contributing to the automaticity of hiPSC-CMs and nodal cardiomyocytes is the substantial presence of the depolarizing funny pacemaker current (If) in combination with no/low levels of IK1, an inwardly rectifier potassium current which stabilizes resting membrane potential of adult human primary atrial and ventricular cardiomyocytes (Goversen et al., 2018; Verkerk et al., 2007). Therefore, we assessed the expression of ion channels that are responsible for If (HCN4, Hyperpolarization Activated Cyclic Nucleotide Gated Potassium Channel 4,) and IK1 (KCNJ2: Kir 2.1, Potassium Channel, Inwardly Rectifying Subfamily J, Member 2 and KCNJ4: Kir 2.3, Potassium Channel, Inwardly Rectifying Subfamily J, Member 4) currents.

Primary cardiomyocyte electrophysiology data show that If is substantially present in nodal cardiomyocytes, low in atrial cardiomyocytes and absent in ventricular cardiomyocytes, and HCN expression is high in nodal, medium in atrial cardiomyocytes and low/absent in human adult ventricular cardiomyocytes (see review by Kane & Terracciano, 2017 As HCN4 expression is linked to pacemaker physiology, it would be an obvious conclusion that HCN4 could be used as nodal marker, for example to select nodal-like hiPSC-CMs. However, our results indicate that HCN4 expression on transcriptomic level is not specific for nodal cardiomyocytes. Interestingly, we found that almost every single cardiomyocyte investigated is positive for HCN4, including human primary ventricular and atrial cardiomyocytes. In line with that, a previous publication (Yechikov et al., 2016) showed that HCN4 is expressed on protein level in 90-92% of the single hiPSC-CMs investigated: Although immunostaining of HCN4 was initially higher in cardiomyocytes showing nodal-like action potentials, differences disappeared with time in culture and downregulation of HCN4. Other publications report an If current in iCell cardiomyocytes (Ma et al., 2011) and in the majority of primary human ventricular cardiomyocytes (Cerbai et al., 2001). Regarding mean ΔCT values, significant differences are found between the five cell groups assessed in the current investigation, with human primary ventricular cardiomyocytes showing the highest mean ΔCT (lowest expression), followed by primary atrial cardiomyocytes, ventricular Pluricytes, iCell cardiomyocytes and atrial Pluricytes (lowest mean ΔCT/highest expression). In summary, ΔCT values for primary human ventricular and atrial cardiomyocytes were higher than for hiPSC-CMs, indicating a lower mean expression level for primary human cardiomyocytes. Besides HCN4, we investigated the expression of KCNJ2 (Kir 2.1) and KCNJ4 (Kir2.3), the ion channel subunits responsible for the conduction of the I_K1_ current. Whether KCNJ2 is the predominant form in primary ventricular cardiomyocytes and KCNJ4 expression is higher in primary atrial cardiomyocytes (Bartos et al., 2015), or whether there are similar expression levels of KCNJ2 and KCNJ4 in atrium and ventricle (Ördög et al., 2006) is reported inconsistently in literature. We observed the following trends in our results: KCNJ2 shows a higher expression (regarding percentage of positive cells and ΔCT) in primary ventricular cardiomyocytes than in primary atrial cardiomyocytes (significantly higher mean ΔCT than primary ventricular cardiomyocytes, atrial and ventricular Pluricytes). This trend also holds true for Pluricytes: While mean ΔCT is comparable between ventricular and atrial Pluricytes, the percentage of positive cells is higher (93%) in ventricular Pluricytes than in atrial Pluricytes (58%). iCell cardiomyocytes show, compared to the other two hiPSC-CM groups, a similar ΔCT value but the lowest percentage of positive cells. KCNJ4 transcripts are detected at similar levels in a majority of primary ventricular and atrial cardiomyocytes. ΔCT values are comparable between the cell groups investigated except for the iCell cardiomyocytes which show a significantly lower mean ΔCT (higher expression, in line to the functional presence of I_K1_ in iCell cardiomyocytes (Ma et al., 2011)). However, the percentage of positive cells in hiPSC-CMs is lower than in primary cardiomyocytes, with the lowest number of positive cells in atrial Pluricytes. Our findings agree with a reported higher expression of KCNJ2 and KCNJ4 in ventricular hiPSC-CMs than in atrial hiPSC-CMs (Cyganek et al., 2018). Although a missing/low I_K1_ is assumed to contribute to the automaticity in hiPSC-CMs, we could detect transcripts of KCNJ2 and KCNJ4. It is difficult to estimate the influence of the two parameters (percentage of positive cells and ΔCT) on the total expression level, making it difficult to compare the cardiomyocytes groups. However, the comparable expression of KCNJ2 (regarding percentage of positive cells and ΔCT) between ventricular Pluricytes and primary ventricular cardiomyocytes could lead, together with highest ΔCT for HCN4 (when considering hiPSC-CMs) in ventricular Pluricytes, to a slower spontaneous beat rate compared to beat rates of iCell cardiomyocytes and atrial Pluricytes. This tendency is in accordance with the mechanism for automaticity in hiPSC-CMs explained above.

To gain further insight into the beating behavior of hiPSC-CMs, we focused on single-cell correlations regarding beat rate and on the comparison of beating and non-beating cells. Despite the association of HCN4, KCNJ2, and KCNJ4 with beat rate and inter-beat interval, no correlation between these parameters could be established on the single-cell level; neither did we find differences between beating and non-beating cells regarding these three channels. Notably, in a previous study, HCN4 expression on protein level was found to correlate with beating frequency following 40 days in culture and correlation became weaker following 60 days (Yechikov et al., 2016). Still, for iCell cardiomyocytes, a correlation of beat rate with CACNA1G (inversely correlated) and inter-beat interval with CACNA1D (positive correlation) could be established. In the heart, both channels show higher expression in nodal and atrial cells and have low/no expression in primary ventricular cardiomyocytes (Kane & Terracciano, 2017). Moreover, both channels are assumed to be involved in nodal automaticity (Baig et al., 2011; Mesirca et al., 2014). This indicates that for iCell cardiomyocytes, these nodal/pacemaking-related ion channels may influence beat rate. Moreover, for iCell cardiomyocytes, beat rate correlates inversely with the fast inward current amplitude (which correlates positively with CT of SCN5A and inversely with upstroke velocity), for ventricular Pluricytes beat rate correlates positively with upstroke velocity (which in turn correlates with the fast inward current amplitude and density) and for atrial Pluricytes, beat rate correlates significantly and inversely with the CT of SCN5A (which correlates inversely with the fast inward current amplitude). These correlations may indicate a role of the fast inward current (sodium current) for pacemaking in hiPSC-CMs. In primary nodal cardiomyocytes of the central SA node, expression of SCN5A is low/absent (Chandler et al., 2009; Kane & Terracciano, 2017) and therefore I_Na_ cannot contribute to pacemaking in these cells. In addition, we observed that beating ventricular Pluricytes have a significantly larger fast inward current amplitude and density and a steeper upstroke velocity (these trends are also seen in iCell cardiomyocytes). In line with these findings, a previous study reported a prominent I_Na_ current in single hESC-derived cardiomyocytes, the inhibition of spontaneous contractions by TTX and concluded that I_Na_/Na_V_1.5 has a prominent role in the generation of spontaneous action potentials in these cells (Satin et al., 2004). Apart from that, comparing beating and non-beating cells within one cell group revealed only few differences in transcriptional and electrophysiological parameters. While some differences do not show an obvious relation to pacemaking mechanisms or to nodal attributes, others would comply with the hypothesis that beating hiPSC-CMs are more nodal-like because the spontaneous contractions are an obvious evidence for the celĺs ability to spontaneously generate action potentials (e.g., higher percentage of CACNA1D- and CACNA1G-positive iCell cardiomyocytes and atrial Pluricytes in beating compared to non-beating cells). Some correlations are at odds with this hypothesis (e.g., non-beating iCell cardiomyocytes show a significantly smaller CT of CACNA1D value (higher expression), a smaller capacitance and shorter APD90 than their beating counterpart).

In summary, our investigation regarding beating behavior and pacemaking of hiPSC-CMs revealed a complex picture and interpretation of results is challenging. On the one hand pacemaking per se is a complex process, presumably involving not only If and I_K1_ (e.g. the so-called calcium clock (Lakatta, Maltsev, & Vinogradova, 2010)). On the other hand, investigation of beating behavior of hiPSC-CMs is complicated by the variable beating behavior of single hiPSC-CMs. We even observed that hiPSC-CMs switched between beating and non-beating states. Such behavior may indicate that pacemaking in hiPSC-CMs is a highly complex process that involves not only the expression of certain ion channel subunit transcripts but also other mechanisms on several levels (e.g. post-transcriptional, post-translational, structural, metabolic and other mechanisms). It is important to note that the beat rate determined in the current study only reflects a snapshot of the beating behavior prior to patch clamp recordings. If absence of automaticity was temporary, no transcriptional differences would be expected. Moreover, we also found non-beating hiPSC-CMs generating spontaneous action potentials in our study. The fact that non-beating cells can produce spontaneous action potentials is an indication that mechanical processes also needed to be considered. A high percentage of hiPSC-CMs has the ability to spontaneously generate action potentials, which only becomes visible for those hiPSC-CMs in which an action potential is followed by a contraction. In some cells and/or in certain periods an action potential may not lead to a contraction. The fact that the contractile apparatus of hiPSC-CMs differs from that of primary adult human cardiomyocytes (Karakikes et al., 2015) may contribute to this observation.

#### 4.3.3 Action potential duration and related ion channel expression (CACNA1C and KCNH2)

In addition to capacitance and spontaneous beat rate, we also determined the action potential duration of single cardiomyocytes. We found that the mean APD90 of iCell cardiomyocytes (219 ms) and ventricular Pluricytes (271 ms) are comparable while the mean APD90 of atrial Pluricytes (40 ms) and primary ventricular cardiomyocytes (110 ms) were significantly shorter. Shorter action potential durations of atrial hiPSC-CMs compared to ventricular hiPSC-CMs were reported in previous studies (Cyganek et al., 2018). When compared to published data of primary atrial cardiomyocytes (200/400 ms; (Kane & Terracciano, 2017)), the mean APD90 of atrial Pluricytes in our study is considered smaller. Moreover, the mean APD90 for iCell cardiomyocytes and ventricular Pluricytes are in the range of previously reported hiPSC-CM APD90 data (123 – 548 ms; (Barbuti et al., 2016). Nevertheless, also the mean APD90 of iCell cardiomyocytes reported in our study is shorter than the APD90 of iCell cardiomyocytes reported previously (Barbuti et al., 2016; Hortigon-Vinagre et al., 2016). This trend continues in primary ventricular cardiomyocytes: The mean APD90 in our study (110 ms) is shorter than reviewed in the Kane et al publication (250/440 ms, see (Kane & Terracciano, 2017)). Another publication also reported slightly shorter APDs for atrial (150 ms) and ventricular (172 ms) tissue (Goodrow et al., 2018). In addition to the morphological heterogeneity of single hiPSC-CMs, we also observed a huge variation in action potential duration of single hiPSC-CMs (see Figure 1, Table 2). Interestingly, the standard deviation of APD90 was also high for primary ventricular cardiomyocytes. While very long (>500 ms) action potentials were only recorded with iCell cardiomyocytes and ventricular Pluricytes, very short (<50 ms) action potentials were recorded for every cell type we investigated. Overall, we found shorter mean action potential durations and higher variations compared to literature. The relatively high percentage of very short action potentials (cells had a healthy visual appearance and no obviously shifted resting potential) lowered mean APD90 and increased heterogeneity. There are many possible reasons for short action potentials. One may be opening of KATP channels under metabolic stress / ATP decline (Faivre & Findlay, 1990; Findlay, Deroubaix, Guiraudou, & Coraboeuf, 1989). Action potentials in the present study were measured right after opening the cell to avoid mRNA loss. However, it is a common procedure in pharmacological experiments to allow the action potential to stabilize before initiation of recording. Our approach may have been too fast to allow ATP to diffuse from the pipette internal solution into the cell, to close KATP channels, and to level out inter-cell variability. Such shortcoming of our approach may explain the discrepancy between our results and published data. Other possible explanations for the observed heterogeneity may be the usage of singularized cells (it has been shown that density of cells can affect AP morphology and duration (Du et al., 2015)) and that we did not preselect cells to perform recordings. Exploration of computational models of action potential may help to get mechanistic insight into ionic currents involved in very short action potentials.

Since duration of the action potential can be influenced with pharmacological modulation of L-type calcium (I_CaL_) and hERG (I_Kr_) currents (Rast et al., 2016), we investigated whether APD90 differences between the cell types and the heterogeneity of single cells within one cell type could be explained by different expression of CACNA1C (the gene for the L-type calcium current I_CaL_ conducting subunit CaV1.2) and/or KCNH2 (the gene for the I_Kr_ conducting subunit K_V_11.1, hERG). In agreement with published data (Kane & Terracciano, 2017), we found almost every single cardiomyocyte to be positive for CACNA1C and KCNH2. Primary cardiomyocytes show higher ΔCT values (lower expression) than hiPSC-CMs. This agrees with published bulk RT-qPCR data of iCell cardiomyocytes displaying higher/similar expression levels for KCNH2 and CACNA1C respectively, than the human heart (Kodama et al., 2019). Moreover, it was reported that I_Kr_ was recorded in every iCell cardiomyocyte investigated (current densities close to values for primary cardiomyocytes) and that current densities of I_CaL_ were larger than for primary human cardiomyocytes (Ma et al., 2011). Despite the significant contribution of I_Kr_ and I_CaL_ to APD90, correlation analysis didńt show a relationship between APD90 and CACNA1C or KCNH2 on the single-cell level. This lack of correlation can be due to the complex regulation of ion channels. For example, I_CaL_ function is known to be regulated by calcium- and voltage-dependent inactivation (Lee, Marban, & Tsien, 1985) and protein kinase A (Pallien & Klussmann, 2020), indicating that regulation of ion currents is a highly complex process which cannot be simplified by assessment of the transcription of ion channel subunits. Moreover, the duration of an action potential is influenced by many other ionic currents and transcripts of just two ion channels may not be sufficient to explain this parameter.

#### 4.3.4 Upstroke velocity, fast inward current amplitude/density and related ion channel expression (SCN5A)

Next, we investigated the upstroke velocity of the action potential, the fast inward current amplitude and its density. A mean upstroke velocity of 310 V/s for primary ventricular cardiomyocytes is in line with reported data (200-300 V/s according to (Kane & Terracciano, 2017)), while our values for hiPSC-CMs (Ventricular Pluricytes (404 V/s), iCell cardiomyocytes (340 V/s) and atrial Pluricytes (208 V/s)) are higher than those reported previously (9-115 V/s according to (Barbuti et al., 2016)). The fast inward current was detected in every cell we investigated. Atrial Pluricytes (-7.55 nA) and primary ventricular cardiomyocytes (-10.11 nA) showed significantly smaller mean amplitudes than iCell cardiomyocytes (-16.12 nA) and ventricular Pluricytes (-17.17 nA). As the hiPSC-CMs show smaller capacitance values than the primary ventricular cardiomyocytes, hiPSC-CMs show significantly higher (iCells: -700.48, ventricular Pluricytes: -791.36) and significantly smaller (atrial Pluricytes with -447.24 pA/pF) fast inward current densities than the primary ventricular cardiomyocytes (-105.32 pA/pF). Comparisons to other studies are difficult because of varying external solutions and electrophysiological protocols. Notably, our values are higher than in other reports (according to reviewed data in (Barbuti et al., 2016): 12.2 -310 pA/pF), maybe due to our negative holding potential (-100 mV) and the physiological sodium concentration (137 mM). Taken this into consideration, we estimate that currents we recorded in the current study are rather under- than over-estimated (technical limitations due to the amplifier technology). Additionally, high standard deviation values occurred in our study can be due to performance of recordings without prior selection of cells. In a previous study, a I_Na_ density of -216.7 pA/pF was recorded in iCell cardiomyocytes, which may be underestimated due to measurements in small cells (mean capacitance of 15.8 pF) and under reduced sodium concentration (Ma et al., 2011). Moreover, a previous study also reports I_Na_ in almost every hiPSC-CM (93%) (Malan, Friedrichs, Fleischmann, & Sasse, 2011). In line with finding the fast inward current in every cell, every cell investigated was positive for SCNA5 (Sodium Voltage-Gated Channel Alpha Subunit 5, NaV1.5 conducting I_Na_). The ΔCT values were comparable between the cell groups except for one pair: Ventricular Pluricytes showed a significantly lower mean ΔCT (higher expression) than primary ventricular cardiomyocytes. In line with that, literature also reports high and comparable expression of SCN5A in atrium and ventricle (Kane & Terracciano, 2017; Ördög et al., 2006) and another study showed comparable SCN5A expression between the human heart and iCell cardiomyocytes in bulk RT-qPCR experiments (Kodama et al., 2019). The reliability of our findings is strengthened by the positive correlation of SCN5A with the fast inward current amplitude and by the inverse correlation of the fast inward current amplitude and density with the upstroke velocity of the action potential for all three hiPSC-CM groups. Thus, in this case, ion channel subunit transcripts seem to translate to electrophysiological measurements. In line with these findings, another study reported a TTX-dependent upstroke of the action potential in iCell cardiomyocytes (Ma et al., 2011).

#### 4.3.5 Ion channel expression associated with primary cardiac subtypes (CACNA1D, CACNA1G, KCNA5)

The ion channel transcripts investigated using the single-cell RT-qPCR approach were chosen due to estimated functional relations to electrophysiological parameters and due to assumed differential expression in primary cardiac subtypes. Different types of cardiomyocytes exist in the adult heart (nodal, atrial and ventricular cells), assuming different function, gene expression, morphology and electrophysiology. Commonly accepted attributes of the prototypes that are of interest for us are depicted in figure 1 in (Kane & Terracciano, 2017). For example, CACNA1D (Calcium Voltage-Gated Channel Subunit Alpha1 D, CaV1.3 conducting I_CaL_) and CACNA1G (Calcium Voltage-Gated Channel Subunit Alpha1 G, CaV3.1, conducting ICaT) are low/not expressed in primary ventricular cardiomyocytes, but show higher expression in nodal and atrial cells (Kane & Terracciano, 2017). Moreover, KCNA5 (Potassium Voltage-Gated Channel Subfamily A Member 5, K_V_1.5, conducting I_Kur_) shows high expression in atrial cardiomyocytes, but it is associated with low expression/absence in the other cardiac cell types (Kane & Terracciano, 2017). Thus, we compared the expression of these three channels in ventricular/atrial Pluricytes and ventricular/atrial primary cardiomyocytes. Our data show that human atrial primary cardiomyocytes and atrial Pluricytes demonstrated a higher percentage of positive cells for CACNA1D and CACNA1G than ventricular primary cardiomyocytes/ventricular Pluricytes, respectively. In fact, mean ΔCT values of CACNA1G and CACNA1D are smaller in the three hiPSC-CM groups, indicating a higher expression of these pacemaking-related ion channels (Baig et al., 2011; Mesirca et al., 2014) compared to the two primary cell groups. In line with literature (Kane & Terracciano, 2017), KCNA5 shows highest expression in atrial primary cardiomyocytes and atrial Pluricytes, having significantly lower mean ΔCT values than every other group and exhibiting a high percentage of positive cells. However, KCNA5 is also expressed in a relatively high percentage of primary ventricular cardiomyocytes but with a higher mean ΔCT, indicating a lower expression level as compared to atrial cardiomyocytes. Other studies also reported the presence of KCNA5 in human ventricular tissue, but with a higher abundance in the atria (Ördög et al., 2006). Moreover, in accordance with our results, the abundance of KCNA5 on the protein level – but without functional current - was reported earlier for primary ventricular cardiomyocytes (Ravens & Cerbai, 2008). Our results for CACNA1G, CACNA1D and KCNA5 in atrial and ventricular Pluricytes reflect a trend for chamber-specificity regarding certain ion channels and highlight the potential of the directed differentiation approach of hiPSC-CMs.

#### 4.3.6 Assessment of cardiac subtypes in hiPSC-CMs

##### 4.3.6.1 Histograms

While ventricular and atrial Pluricytes are thought to represent two different cardiomyocyte subtypes, iCell cardiomyocytes are claimed to be a mixture of nodal-, atrial-, and ventricular-like cardiomyocytes (*iCell Cardiomyocytes Useŕs Guide.*, 2017). Electrophysiological characterization of iCell cardiomyocytes showed that iCell cardiomyocytes display 54% - 64% ventricular-like action potentials, 18% - 22% nodal-like action potentials, and 18% - 24% atrial-like action potentials (Ma et al., 2011; Yonemizu et al., 2019). We hoped to find indications for subpopulations in histograms of expression data. However, this approach turned out not to be suitable, as it did not allow discrimination of subpopulations for pooled primary atrial and ventricular data. An explanation could be that the number of cells needs to be increased in our experiments or that ΔCT values of the investigated targets are not suitable to dissect cardiac subtypes. Further work is needed to clarify this point. Answering the question about the composition of hiPSC-CM populations is of particular interest, since it is still unclear whether three distinct subpopulations exist (nodal-, atrial- and ventricular-like). Although some publications reported three distinct cardiac subtype-related action potential morphologies (*iCell Cardiomyocytes Useŕs Guide.*, 2017; Ma et al., 2011; Yonemizu et al., 2019), other studies indicated that single pluripotent stem cell derived-cardiomyocytes rather show multiple or even a broad spectrum of action potential phenotypes (Ben-Ari et al., 2016; Du et al., 2015; Gorospe et al., 2014; Kane et al., 2016). From a transcriptomic point of view, our failure to dissect distinct hiPSC-CM subpopulations associated with subtypes in a previous single-cell RNA-sequencing study (Schmid et al., 2020) supports the latter conception.

##### 4.3.6.2 Single-cell phenotypes

As distributions of single-cell parameters did not allow conclusions on possible subpopulations in iCell cardiomyocytes, we next focused on the single-cell hiPSC-CM phenotypes (e.g., the combination of electrophysiological and transcriptomic attributes in single cells). A potential and simple outcome of this study could have been to find distinct single-cell phenotypes of the prototypical cardiac subtypes in the iCell cardiomyocyte culture. According to common sense, we would have expected to find nodal-like hiPSC-CMs that beat spontaneously with relatively short action potential and slow upstroke velocity, without a fast inward current and no expression of SCN5A, KCNJ2 or KCNJ4 but being strongly positive for HCN4, CACNA1G and CACNA1D. Moreover, we would have expected to find atrial-like hiPSC-CMs without spontaneous beating, with longer action potential, faster upstroke velocity and a prominent fast inward current and strong expression of KCNA5. In addition, we also expected a prototypical ventricular-like hiPSC-CM to be present in the iCell cardiomyocyte culture with the following characteristics: no spontaneous beating, long action potential, a steep upstroke, prominent fast inward current, no expression of HCN4, CANCA1G and CACNA1D but positive for SCN5A, KCNH2 and KCNJ2. Yet, we could not identify these distinct cardiac-subtype related subpopulations, but we recognized a high percentage of cells with an interesting combination of the investigated parameters. The majority of hiPSC-CMs of all three investigated groups (iCell cardiomyocytes, ventricular and atrial Pluricytes) was beating spontaneously (i.e. able to spontaneously generate action potentials) and was positive for HCN4 (at a higher mean expression level compared to primary atrial or ventricular cardiomyocytes). The same cells also demonstrated a prominent fast inward current that is mainly driven by I_Na_, (SuppFigure 3a) and steep action potential upstroke and were positive for SCN5A. While spontaneous generation of action potentials and a high HCN4 expression are clearly nodal-related pacemaking attributes, the presence of a prominent I_Na_-driven fast inward current, fast upstroke velocity and a high expression of SCN5A are not nodal (but are clearly associated with attributes of adult atrial and ventricular primary cardiomyocytes (Kane & Terracciano, 2017). A previous case report found a large inward current with characteristics of I_Na_ in two out of a total of three human SA node cells analyzed with recording conditions similar to the ones used in the current study (Verkerk, Wilders, van Borren, & Tan, 2009). Considering that expression of SCN5A/NaV1.5 on the transcript/protein level is restricted to the peripheral SA node (Chandler et al., 2009), the two cells in this small sample showing I_Na_-like current might rather have been isolated from the peripheral region of the SA node than from its central region. Thus, an interpretation of the observed single-cell phenotype of a majority of hiPSC-CMs is that these cells combine attributes from different adult cell types.

In summary, our results with hiPSC-CMs not only agree with previously published literature, but the multi-parameter single-cell approach we employed helps us to further complement and strengthen our knowledge of hiPSC-CMs. Our findings correlate well with the characterization of iCell cardiomyocytes model as reported by Ma et al., 2011: Spontaneous beating, heterogeneous action potential morphologies and a TTX-sensitive (and therefore Na-driven) upstroke. In contrast to our interpretation, Ma et al. concluded that iCell cardiomyocytes represent immature cells (which retained pacemaking properties) with nodal-, atrial- and ventricular-like action potentials. The expression of HCN4 in hiPSC-CMs may be seen as immature feature (Später et al., 2013). Since we did not evaluate primary human embryonic cardiomyocytes and assessment of changes in hiPSC-CM characteristics over time is outside the scope of the current study, we cannot conclude on the developmental status of the cells we investigated. Nevertheless, the question of whether hiPSC-CMs can either be seen as immature or neither resemble adult nor fetal primary cardiomyocytes remains to be resolved (e.g., differences in global gene expression has been reported between human embryonic stem cell – derived cardiomyocytes/hiPSC-CMs and both, fetal and adult cardiomyocytes (Gupta et al., 2010)).

#### 4.3.7 Implications for use and final remarks

The fact that the majority of hiPSC-CMs combine nodal and non-nodal attributes may lead to ambiguities with the interpretation of electrophysiological experiments, especially with non-paced cells. For example, it has been published that hiPSC-CMs show an interdependence of repolarization and spontaneous beat rate in MEA experiments (Rast et al., 2016). One potential contributing factor to this phenomenon suggested by the authors of the study is that hiPSC-CMs may have mixed attributes of nodal and ventricular cardiomyocytes. Our results support the hypothesis that the majority of hiPSC-CMs exhibit a mixed cardiac phenotype (combination of nodal and non-nodal attributes) as opposed to have distinct nodal and ventricular cells in the culture which would have been an alternative hypothesis.

Despite very long action potentials being only recorded in hiPSC-CMs, both hiPSC-CMs (mainly iCell cardiomyocytes and ventricular Pluricytes) and primary ventricular cardiomyocytes showed a prominent variability regarding action potential duration. This heterogeneity may cause problems when using the cells for detection of drug-induced changes on action potential duration in a low-throughput assay (usually resulting in small sample sizes) like manual patch-clamp. In this case, selection criteria (use of cells within a defined APD range) or increase of sample size using a higher throughput platform could help to reliably detect changes in action potential duration despite the high basal heterogeneity.

Effects of research substances on individual cardiac ionic currents are typically evaluated in heterologous expression systems. To make an assessment in a more “realistic” physiological environment than in an immortalized cell line, hiPSC-CMs might also be of interest for the analysis of individual cardiac ion currents. As every single hiPSC-CM investigated in this study showed a fast inward current and expression of SCN5A, they are considered a suitable model system for evaluation of effects on the NaV1.5 target. The strong expression of CACNA1C and KCNH2/hERG in almost every cell also indicates that iCell cardiomyocytes and Pluricytes can be considered good model systems for these channels.

The situation is more difficult when the parameter of interest is likely to depend on the integration of multiple ionic currents, like the beat rate and APD. When comparing such integrative parameters, the expression pattern of all relevant ion channels should be comparable, e.g. between hiPSC-CMs and primary cardiomyocytes. While the distributions of SCN5A and KCNH2 seem indeed comparable, a clear shift to the left (higher expression) can be observed for the pacemaking-associated ion channels HCN4, CACNA1G and CACNA1D (see SuppFigure 1). This indicates that the spontaneous depolarization, which impacts the beat rate, upstroke of the action potential and therefore indirectly the action potential duration of some individual hiPSC-CMs might be driven by a combination of If (HCN4), different calcium currents (CACNA1C, CACNA1D, CACNA1G) and I_Na_ (SCN5A). This combination would not be expected in primary central SA nodal, atrial or ventricular cells (Kane & Terracciano, 2017). Therefore, drug-induced changes of integrative parameters, need to be interpreted with the caveat that a set of ion channels different from the expected physiological one might contribute to these parameters in the hiPSC-CM model system being used. A previous case report even describes the presence of a non-cardiac ion channel current in individual batches of hiPSC-CMs (Horváth et al., 2020). Thus, for the analysis of integrative parameters, primary cardiomyocytes might be preferable.

At the beginning of the experiments for this study, our working hypothesis was that a binary presentation of single-cell expression data (Figure 3) would reveal clear, homogeneous and distinguishable subtype-associated expression patterns in primary atrial and ventricular cardiomyocytes. Such presentation should have allowed to derive conclusions on the composition of hiPSC-CM populations. However, we observed different single-cell expression patterns and both positive and negative cells were seen with every target except SCN5A and KCNH2 within one cell group. This points out a certain level of heterogeneity within the five investigated cardiac cell groups, also within primary atrial and ventricular cardiomyocytes, maybe indicating that in addition to prototypical primary nodal, atrial, and ventricular cells, a broad spectrum of overlapping cardiac cell phenotypes can be found in the heart (Kane & Terracciano, 2017). Moreover, although we expected subtype-specific expression of HCN4 (see section 4.4.2) and KCNA5 (see section 4.4.5), our single-cell data indicate that the discrimination of primary atrial and ventricular cardiomyocytes in a binary transcriptomic analysis for these two ion channels is not as clear-cut as expected. Taken together, the ion channel targets we assessed in the current study are not specific for a certain subtype in a (binary) transcriptomic fashion and therefore are not suitable as markers to identify a specific cardiac subtype in a hiPSC-CM culture (if it exists at all – see section 4.4.6.1).

However, the average expression on a population level (Table 3, Figure 4) shows a trend for chamber-specificity regarding certain ion channels, like CACNA1D, CACNA1G, KCNA5 and KCNJ2 for atrial and ventricular Pluricytes when compared to their expression in primary atrial and ventricular cardiomyocytes. Such trend indicates the potential to derive more chamber-like hiPSC-CMs using subtype-directed differentiation approaches. Thus, Pluricytes’ cardiomyocytes may be better suitable to assess chamber-specific ion channel pharmacology than iCell cardiomyocytes which are prepared with an undirected differentiation method.

In conclusion, we assessed hiPSC-CMs for their electrophysiological properties and ion channel expression in order to clarify their utility in different scenarios of safety assessment of novel drugs. We propose that the utilization of hiPSC-CMs for a specific application has to be complemented with a minimal characterization tailored to this specific application.

## 7 Author contributions

**Christina Schmid:** Conceptualization, Data curation, Formal analysis, Investigation, Methodology, Visualization, Writing; **Najah Abi-Gerges:** Procurement of donor hearts and isolation of primary human cardiomyocytes, Proof-reading; **Dietmar Zellner:** Statistics; **Georg Rast:** Conceptualization, Project administration, Resources, Supervision

## Acknowledgements

The authors would like to thank Paul E. Miller, Anh-Tuan Ton, William Nguyen, Ky Truong and Guy Page for technical and administrative assistance and Michael Leitner for valuable input on the manuscript.

## ABBREVIATIONS

APD90: Action potential duration at 90% repolarization
CACNA1C/D/G: Calcium Voltage-Gated Channel Subunit Alpha1 C/D/G
CaV1.2/1.3: voltage-gated L-type calcium channel subunit alpha 1.2/1.3
CaV3.1: voltage-gated T-type calcium channel subunit alpha 3.1
cDNA: Complementary deoxyribonucleic acid
CiPA: Comprehensive *in Vitro* Proarrhythmia Assay
DNA: Deoxyribonucleic acid
CT: Cycle threshold
ΔCT: Normalized cycle threshold
D-PBS: Dulbeccós phosphate buffered saline
EC solution: Extracellular solution
GAPDH: Glycerinaldehyd-3-phosphat-Dehydrogenase
HCN4: Hyperpolarization Activated Cyclic Nucleotide Gated Potassium Channel 4
HERG: human Ether-a-go-go Related Gene
hiPSC-CM: Human induced pluripotent stem cell derived cardiomyocytes
IC solution: Intracellular solution
I_CaL_: L-type (long lasting) calcium current
I_CaT_: T-type (transient) calcium current
I_f_: Funny current
I_K1_: Inward rectifier potassium current
I_Kr_: Rapid component of the delayed rectifier potassium current
I_Kur_: Ultra rapid delayed rectifier potassium current
I_Na_: Cardiac sodium current
KCNA5: Potassium Voltage-Gated Channel Subfamily A Member 5
KCNH2: Potassium Voltage-Gated Channel Subfamily H Member 2
KCNJ2/4: Potassium Channel, Inwardly Rectifying Subfamily J, Member 2/4
Kir2.1/2.3: Inward-rectifier potassium ion channel subunit 2.1/2.3
K_V_ 1.5/ 1.11: voltage gated potassium channel subunit 1.5/ 1.11
mRNA: Messenger ribonucleic acid
RNA: Ribonucleic acid
RNase: Ribonuclease
rt: Room temperature
RT: Reverse transcription
SA: Sino-atrial
SCN5A: Sodium Voltage-Gated Channel Alpha Subunit 5
SD: Standard deviation
single-cell RT-qPCR: Single-cell reverse transcription quantitative polymerase chain reaction
TNNT2: Cardiac muscle troponin T
TTX: Tetrodotoxin

