## Supplemental Material for "Ion channel expression and electrophysiology of singular human (primary and induced pluripotent stem cell derived) cardiomyocytes"

### 1 Supplementary material

#### 1.1 Supplementary figures

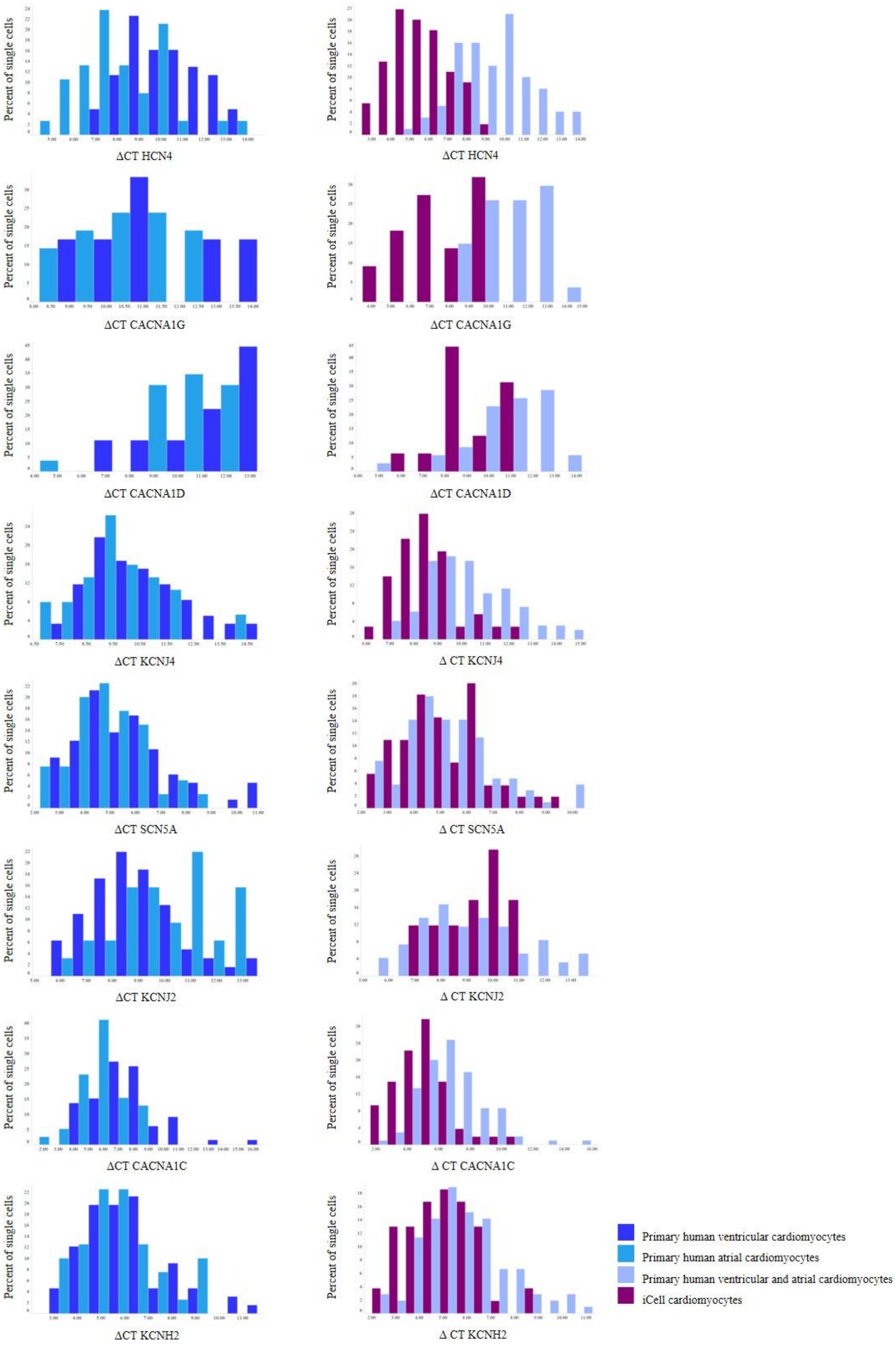

**SuppFigure 1: Histograms of expression parameters.** Comparisons of distributions of single-cell expression data (ΔCT) between atrial and ventricular human primary cardiomyocytes (right side) and between pooled atrial and ventricular primary cardiomyocytes and iCell cardiomyocytes (left side). **Left side**: While for some histograms the distributions are comparable, the expression of HCN4 is left-shifted (higher expression) in primary human atrial cardiomyocytes (note that *n* is low for CACNA1D and CACNA1G). **Right side:** Comparing distributions of single-cell expression data (ΔCT) between pooled atrial and ventricular primary cardiomyocytes and iCell cardiomyocytes shows a slightly smaller bandwidth for iCell cardiomyocytes for some ion channels. While especially the distributions of SCN5A and KCNH2 are comparable, a clear shift to the left (higher expression) can be observed for the pacemaking-associated ion channels HCN4, CACNA1G and CACNA1D**.**

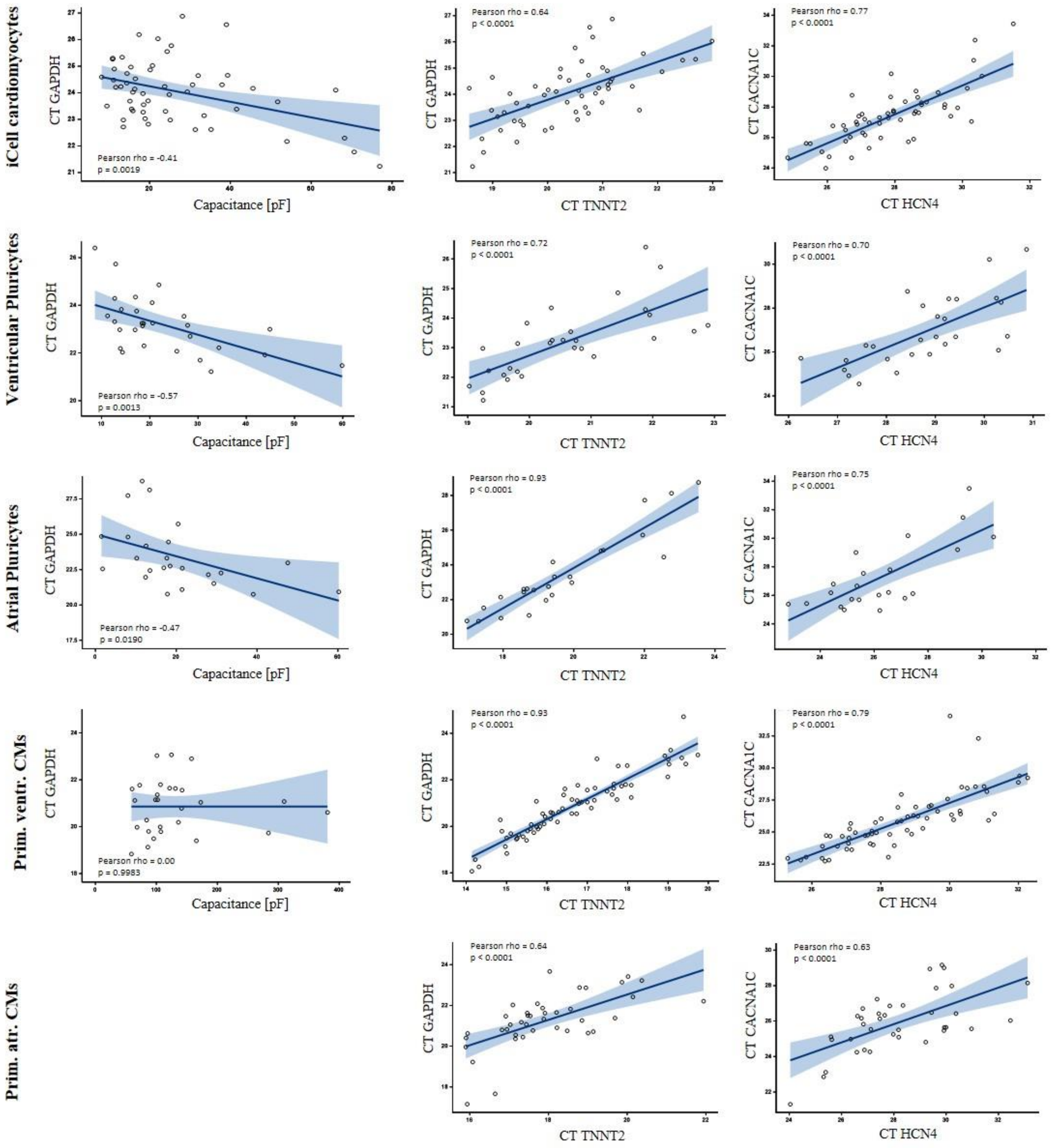


**SuppFigure 2: Correlations of selected parameter pairs.** For hiPSC-CMs, several expression parameters correlate significantly inversely with capacitance (correlations of capacitance with CT of GAPDH are shown exemplary on the left). Moreover, for every cell group, correlations of expression parameters (CT values) with each other revealed strong and highly significant, positive correlations for many parameter pairs (for example, correlations of GAPDH and TNNT2, and CACNA1C and HCN4 are depicted in the middle and on the right). Blue shaded area indicates 95% confidence limits

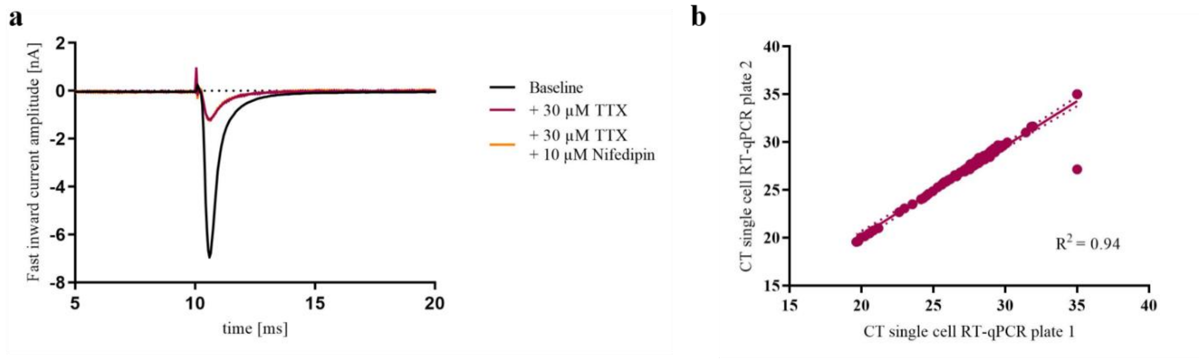


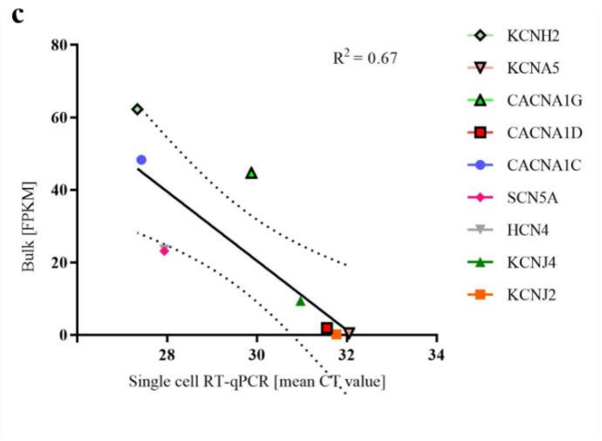


**d**

| | | SCN5A CT Rep. 1 | SCN5A CT Rep. 2 | SCN5A ø CT | CACNA1C CT Rep. 1 | CACNA1C CT Rep. 2 | CACNA1C ø CT |
|---|---|---|---|---|---|---|---|
| **iCell 1** | 20x | 28.8 | 29.0 | 28.9 | 30.1 | 30.1 | 30.1 |
| | 15x | 28.2 | 28.2 | 28.2 | 29.5 | 29.3 | 29.4 |
| | 10x | 27.6 | 27.6 | 27.6 | 28.9 | 28.8 | 28.9 |
| **iCell 2** | 20x | 30.0 | 30.2 | 30.1 | 32.0 | 31.9 | 31.9 |
| | 15x | 29.5 | 29.7 | 29.6 | 31.4 | 31.3 | 31.3 |
| | 10x | 28.8 | 28.8 | 28.8 | 30.6 | 30.8 | 30.7 |

**SuppFigure 3: Reliability and reproducibility of experiments.** All depicted validations have been performed using iCell cardiomyocytes. **(a)** Representative trace of the fast inward current. The current can be blocked with TTX, indicating the cardiac sodium channel $Na_V1.5$ (SCN5A) mainly conducts it. **(b)** The qPCR for different targets (HCN4, SCN5A, CACNA1C, KCNH2, TNNT2 and GAPDH) of 12 iCell cardiomyocytes was run twice on two different plates. CT values of both plates for each cell and target were plotted against each other. High correlation indicates low inter-plate variability. **(c)** Correlation of results of bulk sequencing and mean CT values of iCell cardiomyocytes. **(d)** Representative raw data results of qPCR of two iCell cardiomyocytes (iCell 1 and iCell 2) for SCN5A and CACNA1C. Note that dilutions of pre-amplified cDNA (10 – 15 - 20 x) reflect CT values (difference between 20x and 10x is one CT value) and that CT values of technical replicates are highly reproducible.

#### 1.2 Supplementary tables

**Supp. Table 1: Detailed information on donor hearts.** In total four hearts were used. The number of cells per heart, tested in the current study, are listed. Abbreviations: F, Female; M, Male; BMI, Body Mass Index; COD, Cause Of Death; EF, Ejection Fraction; CVA, Cerebrovascular Accident; ICH, Intracranial Haemorrhage; [a]Organ procurement organization could not transplant the heart and consequently no echocardiography was performed; N/A, Not available.

| Heart# | # of atrial cells | # of ventr. cells | Age | Sex | Ethnicity | BMI | COD | EF (%) |
|---|---|---|---|---|---|---|---|---|
| 1 | 20 | 44 | 44 | F | Hispanic | 23.9 | Anoxia | 70 |
| 2 | - | 20 | 50 | F | Caucasian | 27.0 | CVA/ICH/Stroke | 70 |
| 3 | 1 | 26 | 48 | M | Hispanic | 29.7 | CVA/ICH/Stroke | 73 |
| 5 | 19 | - | 48 | M | Caucasian | 25.0 | Head Trauma/Blunt Injury | N/A[a] |

**Supp. Table 2: Single-cell correlations for iCell cardiomyocytes.** For every pairwise correlation, Pearson´s correlation coefficient r, the p-value and *n* is indicated. For expression parameters (CT), only positive cells are included in the analysis. Significant correlations ($p < 0.05$) are highlighted

| Pearson Correlation Coefficients<br>Prob > \|r\| under H0: Rho=0<br>Number of Observations | | | | | | | | | |
|---|---|---|---|---|---|---|---|---|---|
| | Capacitance [pF] | CT TNNT2 | CT GAPDH | Beat rate [bpm] | Inter-beat interval [ms] | Fast inward current amplitude [nA] | Fast inward current density [nA/pF] | Upstroke velocity [V/ms] | APD90 [ms] |
| Capacitance [pF] | | | | | | | | | |
| CT TNNT2 | -0.47948<br>0.0002<br>55 | | | | | | | | |
| CT GAPDH | -0.41380<br>0.0019<br>54 | 0.63821<br><.0001<br>54 | | | | | | | |
| Beat rate [bpm] | 0.39615<br>0.0006<br>72 | -0.20614<br>0.1311<br>55 | -0.09661<br>0.4871<br>54 | | | | | | |
| Inter-beat interval [ms] | -0.25972<br>0.0451<br>60 | 0.08118<br>0.5875<br>47 | 0.08924<br>0.5553<br>46 | -0.86572<br><.0001<br>60 | | | | | |
| Fast inward current amplitude [nA] | -0.52661<br><.0001<br>71 | 0.25793<br>0.0573<br>55 | 0.40036<br>0.0027<br>54 | -0.32099<br>0.0063<br>71 | 0.22271<br>0.0900<br>59 | | | | |
| Fast inward current density [nA/pF] | 0.22838<br>0.0554<br>71 | -0.18581<br>0.1744<br>55 | 0.07108<br>0.6095<br>54 | -0.04794<br>0.6914<br>71 | 0.02049<br>0.8776<br>59 | 0.57352<br><.0001<br>71 | | | |
| Upstroke velocity [V/ms] | 0.08702<br>0.4738<br>70 | -0.09910<br>0.4716<br>55 | 0.04812<br>0.7297<br>54 | 0.16216<br>0.1799<br>70 | -0.16066<br>0.2242<br>59 | -0.41629<br>0.0004<br>69 | -0.46923<br><.0001<br>69 | | |
| APD90 [ms] | 0.08992<br>0.4526<br>72 | -0.05848<br>0.6715<br>55 | 0.09803<br>0.4807<br>54 | -0.14126<br>0.2366<br>72 | 0.37258<br>0.0034<br>60 | 0.05834<br>0.6289<br>71 | 0.08764<br>0.4674<br>71 | 0.18586<br>0.1234<br>70 | |
| CT KCNJ4 | -0.14583<br>0.3961<br>36 | 0.25792<br>0.1288<br>36 | 0.33011<br>0.0528<br>35 | -0.03112<br>0.8570<br>36 | -0.10542<br>0.5793<br>30 | 0.06821<br>0.6926<br>36 | -0.03945<br>0.8193<br>36 | 0.36794<br>0.0273<br>36 | -0.09641<br>0.5759<br>36 |
| CT HCN4 | -0.13913<br>0.3110<br>55 | 0.57387<br><.0001<br>55 | 0.62395<br><.0001<br>54 | -0.12547<br>0.3614<br>55 | 0.16976<br>0.2540<br>47 | 0.11026<br>0.4229<br>55 | 0.01460<br>0.9157<br>55 | -0.01276<br>0.9264<br>55 | 0.11805<br>0.3907<br>55 |
| CT SCN5A | -0.31477<br>0.0193<br>55 | 0.60979<br><.0001<br>55 | 0.57934<br><.0001<br>54 | -0.22416<br>0.0999<br>55 | 0.08396<br>0.5747<br>47 | 0.33424<br>0.0126<br>55 | 0.12547<br>0.3614<br>55 | -0.24820<br>0.0677<br>55 | -0.06381<br>0.6435<br>55 |
| CT CACNA1C | -0.15034<br>0.2779<br>54 | 0.65659<br><.0001<br>54 | 0.54573<br><.0001<br>53 | -0.05731<br>0.6806<br>54 | -0.06909<br>0.6482<br>46 | 0.13894<br>0.3164<br>54 | -0.03184<br>0.8192<br>54 | 0.11060<br>0.4259<br>54 | 0.07072<br>0.6113<br>54 |
| CT KCNH2 | -0.33075<br>0.0146<br>54 | 0.66434<br><.0001<br>54 | 0.67170<br><.0001<br>53 | -0.27414<br>0.0449<br>54 | 0.20608<br>0.1694<br>46 | 0.24597<br>0.0730<br>54 | -0.04697<br>0.7359<br>54 | 0.02913<br>0.8344<br>54 | 0.02126<br>0.8787<br>54 |
| CT CACNA1D | -0.51714<br>0.0402<br>16 | 0.21706<br>0.4194<br>16 | 0.35693<br>0.1747<br>16 | -0.39439<br>0.1306<br>16 | 0.69038<br>0.0044<br>15 | 0.12324<br>0.6493<br>16 | -0.30098<br>0.2573<br>16 | 0.17883<br>0.5075<br>16 | 0.34933<br>0.1848<br>16 |
| CT CACNA1G | -0.62623<br>0.0018<br>22 | 0.26047<br>0.2417<br>22 | 0.50993<br>0.0153<br>22 | -0.52551<br>0.0120<br>22 | 0.31237<br>0.1800<br>20 | 0.30266<br>0.1710<br>22 | -0.09500<br>0.6741<br>22 | 0.04655<br>0.8370<br>22 | 0.34537<br>0.1154<br>22 |
| CT KCNA5 | 0.20682<br>0.4422<br>16 | 0.43548<br>0.0918<br>16 | 0.24729<br>0.3558<br>16 | -0.04316<br>0.8739<br>16 | -0.23740<br>0.4348<br>13 | 0.07185<br>0.7915<br>16 | 0.21550<br>0.4228<br>16 | -0.06697<br>0.8054<br>16 | -0.28791<br>0.2796<br>16 |
| CT KCNJ2 | 0.28982<br>0.2592<br>17 | 0.24953<br>0.3341<br>17 | 0.35331<br>0.1642<br>17 | 0.15514<br>0.5521<br>17 | -0.21029<br>0.4519<br>15 | 0.09899<br>0.7054<br>17 | 0.60831<br>0.0096<br>17 | -0.47493<br>0.0540<br>17 | -0.53779<br>0.0260<br>17 |

| earson Correlation Coefficients<br>Prob > \|r\| under H0: Rho=0<br>Number of Observations | | | | | | | | | |
|---|---|---|---|---|---|---|---|---|---|
| | CT KCNJ4 | CT HCN4 | CT SCN5A | CT CACNA1C | CT KCNH2 | CT CACNA1D | CT CACNA1G | CT KCNA5 | CT KCNJ2 |
| Capacitance [pF] | | | | | | | | | |
| CT TNNT2 | | | | | | | | | |
| CT GAPDH | | | | | | | | | |
| Beat rate [bpm] | | | | | | | | | |
| Inter-beat interval [ms] | | | | | | | | | |
| Fast inward current amplitude [nA] | | | | | | | | | |
| Fast inward current density [nA/pF] | | | | | | | | | |
| Upstroke velocity [V/ms] | | | | | | | | | |
| APD90 [ms] | | | | | | | | | |
| CT KCNJ4 | | | | | | | | | |
| CT HCN4 | 0.46450<br>0.0043<br>36 | | | | | | | | |
| CT SCN5A | 0.28567<br>0.0912<br>36 | 0.63050<br><.0001<br>55 | | | | | | | |
| CT CACNA1C | 0.48559<br>0.0031<br>35 | 0.77101<br><.0001<br>54 | 0.63570<br><.0001<br>54 | | | | | | |
| CT KCNH2 | 0.54151<br>0.0008<br>35 | 0.74212<br><.0001<br>54 | 0.63704<br><.0001<br>54 | 0.63982<br><.0001<br>53 | | | | | |
| CT CACNA1D | 0.26174<br>0.4112<br>12 | 0.38780<br>0.1378<br>16 | 0.27524<br>0.3022<br>16 | 0.20048<br>0.4566<br>16 | 0.45558<br>0.0879<br>15 | | | | |
| CT CACNA1G | 0.35683<br>0.1597<br>17 | 0.34438<br>0.1165<br>22 | 0.42360<br>0.0495<br>22 | 0.33256<br>0.1305<br>22 | 0.46296<br>0.0300<br>22 | 0.72164<br>0.0433<br>8 | | | |
| CT KCNA5 | 0.08566<br>0.8140<br>10 | 0.35294<br>0.1800<br>16 | 0.34730<br>0.1875<br>16 | 0.51606<br>0.0407<br>16 | 0.40845<br>0.1163<br>16 | -0.75064<br>0.0519<br>7 | -0.23409<br>0.6134<br>7 | | |
| CT KCNJ2 | 0.42002<br>0.1349<br>14 | 0.43623<br>0.0800<br>17 | 0.52937<br>0.0289<br>17 | 0.28982<br>0.2592<br>17 | 0.35981<br>0.1560<br>17 | -0.09204<br>0.8004<br>10 | -0.00349<br>0.9935<br>8 | 0.75423<br>0.0501<br>7 | |

**Supp. Table 3: Single-cell correlations for ventricular Pluricytes.** For every pairwise correlation, Pearson´s correlation coefficient r, the p-value and *n* is indicated. For expression parameters (CT), only positive cells are included in the analysis. Significant correlations ($p < 0.05$) are highlighted.

Pearson Correlation Coefficients
Prob > |r| under H0: Rho=0
Number of Observations

| | Capacitance [pF] | CT TNNT2 | CT GAPDH | Beat rate [bpm] | Inter-beat interval [ms] | Fast inward current amplitude [nA] | Fast inward current density [nA/pF] | Upstroke velocity [V/ms] | APD90 [ms] |
|---|---|---|---|---|---|---|---|---|---|
| Capacitance [pF] | | | | | | | | | |
| CT TNNT2 | -0.47201<br>0.0097<br>29 | | | | | | | | |
| CT GAPDH | -0.56797<br>0.0013<br>29 | 0.72151<br><.0001<br>29 | | | | | | | |
| Beat rate [bpm] | 0.25750<br>0.1548<br>32 | 0.22268<br>0.2456<br>29 | 0.14911<br>0.4401<br>29 | | | | | | |
| Inter-beat interval [ms] | -0.31433<br>0.1652<br>21 | -0.11077<br>0.6617<br>18 | 0.05129<br>0.8398<br>18 | -0.87122<br><.0001<br>21 | | | | | |
| Fast inward current amplitude [nA] | -0.55135<br>0.0019<br>29 | 0.33893<br>0.0903<br>26 | 0.25595<br>0.2069<br>26 | -0.28312<br>0.1367<br>29 | 0.05048<br>0.8374<br>19 | | | | |
| Fast inward current density [nA/pF] | 0.38774<br>0.0377<br>29 | -0.24292<br>0.2318<br>26 | -0.40638<br>0.0394<br>26 | -0.09831<br>0.6119<br>29 | -0.20326<br>0.4039<br>19 | 0.48312<br>0.0079<br>29 | | | |
| Upstroke velocity [V/ms] | 0.32201<br>0.0827<br>30 | 0.09515<br>0.6369<br>27 | -0.05456<br>0.7869<br>27 | 0.44634<br>0.0134<br>30 | -0.11209<br>0.6478<br>19 | -0.54228<br>0.0024<br>29 | -0.39389<br>0.0345<br>29 | | |
| APD90 [ms] | 0.14464<br>0.4296<br>32 | -0.32318<br>0.0873<br>29 | -0.33659<br>0.0742<br>29 | -0.08132<br>0.6582<br>32 | 0.37459<br>0.0943<br>21 | 0.03141<br>0.8715<br>29 | 0.20910<br>0.2763<br>29 | 0.19974<br>0.2900<br>30 | |
| CT KCNJ4 | 0.25169<br>0.3137<br>18 | 0.35900<br>0.1434<br>18 | 0.26974<br>0.2790<br>18 | -0.17098<br>0.4975<br>18 | 0.20517<br>0.5451<br>11 | 0.18620<br>0.4899<br>16 | 0.30188<br>0.2558<br>16 | -0.13674<br>0.6008<br>17 | 0.09025<br>0.7217<br>18 |
| CT HCN4 | -0.58489<br>0.0014<br>27 | 0.54664<br>0.0032<br>27 | 0.80464<br><.0001<br>27 | -0.09305<br>0.6443<br>27 | 0.32122<br>0.2087<br>17 | 0.28815<br>0.1721<br>24 | -0.38461<br>0.0635<br>24 | -0.08920<br>0.6716<br>25 | -0.31381<br>0.1109<br>27 |
| CT SCN5A | -0.54420<br>0.0023<br>29 | 0.73648<br><.0001<br>29 | 0.66461<br><.0001<br>29 | -0.14862<br>0.4416<br>29 | 0.08088<br>0.7497<br>18 | 0.54891<br>0.0037<br>26 | -0.04690<br>0.8200<br>26 | -0.29704<br>0.1324<br>27 | -0.17614<br>0.3607<br>29 |
| CT CACNA1C | -0.26417<br>0.1661<br>29 | 0.70566<br><.0001<br>29 | 0.78418<br><.0001<br>29 | 0.17565<br>0.3621<br>29 | 0.17628<br>0.4841<br>18 | 0.11549<br>0.5742<br>26 | -0.29148<br>0.1485<br>26 | 0.24990<br>0.2087<br>27 | -0.28120<br>0.1395<br>29 |
| CT KCNH2 | -0.50268<br>0.0064<br>28 | 0.83824<br><.0001<br>28 | 0.77068<br><.0001<br>28 | 0.18654<br>0.3419<br>28 | 0.00932<br>0.9707<br>18 | 0.41888<br>0.0372<br>25 | -0.07150<br>0.7341<br>25 | -0.09468<br>0.6455<br>26 | -0.28008<br>0.1489<br>28 |
| CT CACNA1D | -0.60008<br>0.1543<br>7 | 0.13146<br>0.7787<br>7 | 0.50153<br>0.2515<br>7 | -0.16058<br>0.7309<br>7 | -0.34070<br>0.6593<br>4 | 0.42476<br>0.3422<br>7 | -0.00830<br>0.9859<br>7 | -0.45459<br>0.3055<br>7 | 0.04534<br>0.9231<br>7 |
| CT CACNA1G | -0.61184<br>0.1443<br>7 | 0.20023<br>0.6668<br>7 | 0.33206<br>0.4668<br>7 | -0.26131<br>0.5714<br>7 | -0.50815<br>0.3820<br>5 | 0.89202<br>0.0069<br>7 | 0.47738<br>0.2787<br>7 | -0.23095<br>0.6183<br>7 | -0.62289<br>0.1351<br>7 |
| CT KCNA5 | 0.05814<br>0.8504<br>13 | 0.05273<br>0.8642<br>13 | 0.15099<br>0.6224<br>13 | 0.38399<br>0.1952<br>13 | -0.27364<br>0.5998<br>6 | -0.48551<br>0.0926<br>13 | -0.55685<br>0.0481<br>13 | 0.51997<br>0.0685<br>13 | -0.54486<br>0.0542<br>13 |
| CT KCNJ2 | -0.15219<br>0.4486<br>27 | 0.53316<br>0.0042<br>27 | 0.46542<br>0.0144<br>27 | -0.08619<br>0.6691<br>27 | 0.16289<br>0.5467<br>16 | 0.08833<br>0.6746<br>25 | -0.20747<br>0.3197<br>25 | -0.09146<br>0.6568<br>26 | -0.04127<br>0.8381<br>27 |

| Pearson Correlation Coefficients<br>Prob > \|r\| under H0: Rho=0<br>Number of Observations | | | | | | | | | |
|---|---|---|---|---|---|---|---|---|---|
| | CT KCNJ4 | CT HCN4 | CT SCN5A | CT CACNA1C | CT KCNH2 | CT CACNA1D | CT CACNA1G | CT KCNA5 | CT KCNJ2 |
| Capacitance [pF] | | | | | | | | | |
| CT TNNT2 | | | | | | | | | |
| CT GAPDH | | | | | | | | | |
| Beat rate [bpm] | | | | | | | | | |
| Inter-beat interval [ms] | | | | | | | | | |
| Fast inward current amplitude [nA] | | | | | | | | | |
| Fast inward current density [nA/pF] | | | | | | | | | |
| Upstroke velocity [V/ms] | | | | | | | | | |
| APD90 [ms] | | | | | | | | | |
| CT KCNJ4 | | | | | | | | | |
| CT HCN4 | 0.26439<br>0.3051<br>17 | | | | | | | | |
| CT SCN5A | 0.31327<br>0.2056<br>18 | 0.58871<br>0.0012<br>27 | | | | | | | |
| CT CACNA1C | 0.39897<br>0.1010<br>18 | 0.69500<br><.0001<br>27 | 0.65481<br>0.0001<br>29 | | | | | | |
| CT KCNH2 | 0.29053<br>0.2579<br>17 | 0.58006<br>0.0019<br>26 | 0.70330<br><.0001<br>28 | 0.64720<br>0.0002<br>28 | | | | | |
| CT CACNA1D | -0.72842<br>0.1006<br>6 | 0.10944<br>0.8153<br>7 | 0.71453<br>0.0712<br>7 | 0.34557<br>0.4477<br>7 | 0.38486<br>0.3939<br>7 | | | | |
| CT CACNA1G | -0.22706<br>0.6653<br>6 | 0.56515<br>0.1861<br>7 | 0.44023<br>0.3229<br>7 | 0.31254<br>0.4950<br>7 | 0.28417<br>0.5368<br>7 | 0.57174<br>0.4283<br>4 | | | |
| CT KCNA5 | -0.70958<br>0.0487<br>8 | -0.00728<br>0.9830<br>11 | -0.39804<br>0.1780<br>13 | 0.19096<br>0.5320<br>13 | 0.06333<br>0.8372<br>13 | -0.97117<br>0.0288<br>4 | -0.96253<br>0.0375<br>4 | | |
| CT KCNJ2 | 0.39873<br>0.1012<br>18 | 0.44569<br>0.0256<br>25 | 0.69364<br><.0001<br>27 | 0.48381<br>0.0106<br>27 | 0.55861<br>0.0030<br>26 | 0.26194<br>0.5704<br>7 | -0.53098<br>0.2201<br>7 | -0.32365<br>0.2807<br>13 | |

**Supp. Table 4: Single-cell correlations for atrial Pluricytes.** For every pairwise correlation, Pearson´s correlation coefficient r, the p-value and *n* is indicated. For expression parameters (CT), only positive cells are included in the analysis. Significant correlations ($p < 0.05$) are highlighted.

| Pearson Correlation Coefficients<br>Prob > \|r\| under H0: Rho=0<br>Number of Observations | | | | | | | | | |
|---|---|---|---|---|---|---|---|---|---|
| | Capacitance [pF] | CT TNNT2 | CT GAPDH | Beat rate [bpm] | Inter-beat interval [ms] | Fast inward current amplitude [nA] | Fast inward current density [nA/pF] | Upstroke velocity [V/ms] | APD90 [ms] |
| Capacitance [pF] | | | | | | | | | |
| CT TNNT2 | -0.40012<br>0.0527<br>24 | | | | | | | | |
| CT GAPDH | -0.47483<br>0.0190<br>24 | 0.92612<br><.0001<br>24 | | | | | | | |
| Beat rate [bpm] | 0.05421<br>0.7645<br>33 | -0.34275<br>0.1011<br>24 | -0.18465<br>0.3877<br>24 | | | | | | |
| Inter-beat interval [ms] | -0.42786<br>0.0676<br>19 | -0.16396<br>0.6106<br>12 | -0.33029<br>0.2944<br>12 | -0.96566<br><.0001<br>19 | | | | | |
| Fast inward current amplitude [nA] | 0.13761<br>0.4937<br>27 | 0.11004<br>0.6638<br>18 | 0.00055<br>0.9983<br>18 | -0.14726<br>0.4636<br>27 | 0.17213<br>0.5089<br>17 | | | | |
| Fast inward current density [nA/pF] | 0.44261<br>0.0208<br>27 | -0.08424<br>0.7396<br>18 | -0.23965<br>0.3382<br>18 | 0.02145<br>0.9154<br>27 | -0.03594<br>0.8911<br>17 | 0.87199<br><.0001<br>27 | | | |
| Upstroke velocity [V/ms] | -0.20676<br>0.3008<br>27 | -0.18299<br>0.4674<br>18 | -0.00297<br>0.9907<br>18 | 0.19704<br>0.3246<br>27 | -0.13948<br>0.5934<br>17 | -0.70504<br><.0001<br>27 | -0.64460<br>0.0003<br>27 | | |
| APD90 [ms] | 0.43673<br>0.0178<br>29 | -0.13310<br>0.5759<br>20 | -0.20454<br>0.3870<br>20 | -0.19823<br>0.3026<br>29 | 0.27963<br>0.2611<br>18 | 0.21728<br>0.2763<br>27 | 0.26705<br>0.1781<br>27 | -0.28646<br>0.1474<br>27 | |
| CT KCNJ4 | 0.04194<br>0.9084<br>10 | 0.41908<br>0.2280<br>10 | 0.38299<br>0.2747<br>10 | -0.25614<br>0.4750<br>10 | -0.39990<br>0.5047<br>5 | 0.81202<br>0.0497<br>6 | 0.73600<br>0.0953<br>6 | -0.44495<br>0.3766<br>6 | 0.01030<br>0.9807<br>8 |
| CT HCN4 | -0.26246<br>0.2153<br>24 | 0.83107<br><.0001<br>24 | 0.82874<br><.0001<br>24 | -0.34095<br>0.1030<br>24 | -0.01425<br>0.9649<br>12 | -0.02507<br>0.9213<br>18 | -0.29049<br>0.2423<br>18 | -0.04718<br>0.8525<br>18 | -0.07585<br>0.7506<br>20 |
| CT SCN5A | -0.10878<br>0.6129<br>24 | 0.59705<br>0.0021<br>24 | 0.48698<br>0.0158<br>24 | -0.42486<br>0.0385<br>24 | 0.45904<br>0.1333<br>12 | 0.58832<br>0.0102<br>18 | 0.36252<br>0.1393<br>18 | -0.35062<br>0.1537<br>18 | 0.29375<br>0.2087<br>20 |
| CT CACNA1C | 0.01258<br>0.9557<br>22 | 0.74915<br><.0001<br>22 | 0.62045<br>0.0021<br>22 | -0.37129<br>0.0889<br>22 | -0.02613<br>0.9357<br>12 | 0.44823<br>0.0816<br>16 | 0.29919<br>0.2603<br>16 | -0.29797<br>0.2623<br>16 | 0.09169<br>0.7175<br>18 |
| CT KCNH2 | -0.32193<br>0.1250<br>24 | 0.91934<br><.0001<br>24 | 0.93871<br><.0001<br>24 | -0.26181<br>0.2165<br>24 | -0.20756<br>0.5174<br>12 | 0.20871<br>0.4059<br>18 | -0.05058<br>0.8420<br>18 | -0.16894<br>0.5028<br>18 | -0.00881<br>0.9706<br>20 |
| CT CACNA1D | -0.02504<br>0.9267<br>16 | 0.24294<br>0.3646<br>16 | 0.29601<br>0.2656<br>16 | -0.33018<br>0.2117<br>16 | 0.04619<br>0.8992<br>10 | -0.35265<br>0.2609<br>12 | -0.55552<br>0.0608<br>12 | 0.02621<br>0.9356<br>12 | 0.25745<br>0.3742<br>14 |
| CT CACNA1G | -0.27952<br>0.3130<br>15 | 0.36712<br>0.1783<br>15 | 0.33911<br>0.2163<br>15 | -0.50443<br>0.0552<br>15 | 0.63183<br>0.0679<br>9 | 0.08375<br>0.7958<br>12 | -0.12437<br>0.7002<br>12 | -0.03069<br>0.9246<br>12 | 0.03060<br>0.9209<br>13 |
| CT KCNA5 | -0.42523<br>0.0431<br>23 | 0.87963<br><.0001<br>23 | 0.74687<br><.0001<br>23 | -0.28490<br>0.1876<br>23 | 0.11243<br>0.7279<br>12 | 0.05977<br>0.8137<br>18 | -0.10873<br>0.6676<br>18 | -0.15860<br>0.5296<br>18 | -0.32212<br>0.1660<br>20 |
| CT KCNJ2 | -0.21663<br>0.4570<br>14 | 0.29410<br>0.3074<br>14 | 0.16978<br>0.5617<br>14 | -0.53080<br>0.0508<br>14 | 0.66032<br>0.0747<br>8 | 0.35243<br>0.2878<br>11 | 0.09561<br>0.7798<br>11 | -0.42227<br>0.1958<br>11 | 0.52114<br>0.0678<br>13 |

| Pearson Correlation Coefficients<br>Prob > \|r\| under H0: Rho=0<br>Number of Observations | | | | | | | | | |
|---|---|---|---|---|---|---|---|---|---|
| | CT KCNJ4 | CT HCN4 | CT SCN5A | CT CACNA1C | CT KCNH2 | CT CACNA1D | CT CACNA1G | CT KCNA5 | CT KCNJ2 |
| Capacitance [pF] | | | | | | | | | |
| CT TNNT2 | | | | | | | | | |
| CT GAPDH | | | | | | | | | |
| Beat rate [bpm] | | | | | | | | | |
| Inter-beat interval [ms] | | | | | | | | | |
| Fast inward current amplitude [nA] | | | | | | | | | |
| Fast inward current density [nA/pF] | | | | | | | | | |
| Upstroke velocity [V/ms] | | | | | | | | | |
| APD90 [ms] | | | | | | | | | |
| CT KCNJ4 | | | | | | | | | |
| CT HCN4 | 0.46375<br>0.1770<br>10 | | | | | | | | |
| CT SCN5A | 0.22506<br>0.5319<br>10 | 0.63971<br>0.0008<br>24 | | | | | | | |
| CT CACNA1C | 0.45614<br>0.1852<br>10 | 0.74993<br><.0001<br>22 | 0.74429<br><.0001<br>22 | | | | | | |
| CT KCNH2 | 0.59746<br>0.0682<br>10 | 0.87736<br><.0001<br>24 | 0.69341<br>0.0002<br>24 | 0.78575<br><.0001<br>22 | | | | | |
| CT CACNA1D | -0.26572<br>0.4895<br>9 | 0.54085<br>0.0305<br>16 | 0.22235<br>0.4078<br>16 | 0.21266<br>0.4291<br>16 | 0.33062<br>0.2110<br>16 | | | | |
| CT CACNA1G | 0.03913<br>0.9267<br>8 | 0.27949<br>0.3130<br>15 | 0.58176<br>0.0229<br>15 | 0.50953<br>0.0524<br>15 | 0.32717<br>0.2339<br>15 | 0.02963<br>0.9235<br>13 | | | |
| CT KCNA5 | 0.03325<br>0.9273<br>10 | 0.72943<br><.0001<br>23 | 0.58483<br>0.0034<br>23 | 0.70372<br>0.0004<br>21 | 0.73649<br><.0001<br>23 | -0.13819<br>0.6098<br>16 | 0.48246<br>0.0685<br>15 | | |
| CT KCNJ2 | -0.65207<br>0.1125<br>7 | -0.07961<br>0.7868<br>14 | 0.48736<br>0.0771<br>14 | 0.20556<br>0.5005<br>13 | 0.24389<br>0.4007<br>14 | 0.63635<br>0.0353<br>11 | 0.35503<br>0.2840<br>11 | 0.04165<br>0.8876<br>14 | |

**Supp. Table 5: Single-cell correlations for primary human ventricular cardiomyocytes.** For every pairwise correlation, Pearson´s correlation coefficient r, the p-value and *n* is indicated. For expression parameters (CT), only positive cells are included in the analysis. Significant correlations ($p < 0.05$) are highlighted.

| Pearson Correlation Coefficients<br>Prob > \|r\| under H0: Rho=0<br>Number of Observations | | | | | | | | | |
|---|---|---|---|---|---|---|---|---|---|
| | Capacitance [pF] | CT TNNT2 | CT GAPDH | Beat rate [bpm] | Inter-beat interval [ms] | Fast inward current amplitude [nA] | Fast inward current density [nA/pF] | Upstroke velocity [V/ms] | APD90 [ms] |
| Capacitance [pF] | | | | | | | | | |
| CT TNNT2 | -0.07571<br>0.7018<br>28 | | | | | | | | |
| CT GAPDH | 0.00042<br>0.9983<br>28 | 0.92943<br><.0001<br>66 | | | | | | | |
| Beat rate [bpm] | | | | | | | | | |
| Inter-beat interval [ms] | | | | | | | | | |
| Fast inward current amplitude [nA] | -0.02630<br>0.8639<br>45 | -0.23795<br>0.2227<br>28 | -0.10766<br>0.5856<br>28 | | | | | | |
| Fast inward current density [nA/pF] | 0.59838<br><.0001<br>45 | -0.12327<br>0.5320<br>28 | 0.00335<br>0.9865<br>28 | | | 0.59377<br><.0001<br>45 | | | |
| Upstroke velocity [V/ms] | -0.13940<br>0.4625<br>30 | 0.15464<br>0.5273<br>19 | 0.15104<br>0.5371<br>19 | | | -0.10086<br>0.5893<br>31 | -0.17327<br>0.3598<br>30 | | |
| APD90 [ms] | 0.27787<br>0.0646<br>45 | -0.22441<br>0.2419<br>29 | -0.33484<br>0.0758<br>29 | | | -0.22486<br>0.1330<br>46 | -0.04896<br>0.7494<br>45 | 0.08886<br>0.6173<br>34 | |
| CT KCNJ4 | -0.17585<br>0.3803<br>27 | 0.68812<br><.0001<br>60 | 0.53555<br><.0001<br>60 | | | -0.23392<br>0.2403<br>27 | -0.14675<br>0.4651<br>27 | 0.33120<br>0.1794<br>18 | -0.10184<br>0.6061<br>28 |
| CT HCN4 | 0.07562<br>0.7021<br>28 | 0.83212<br><.0001<br>62 | 0.79769<br><.0001<br>62 | | | -0.29168<br>0.1321<br>28 | -0.03873<br>0.8449<br>28 | 0.35509<br>0.1357<br>19 | -0.22993<br>0.2302<br>29 |
| CT SCN5A | -0.09626<br>0.6261<br>28 | 0.93413<br><.0001<br>66 | 0.86351<br><.0001<br>66 | | | -0.33872<br>0.0779<br>28 | -0.17193<br>0.3817<br>28 | 0.21159<br>0.3845<br>19 | -0.15483<br>0.4226<br>29 |
| CT CACNA1C | 0.07060<br>0.7211<br>28 | 0.85971<br><.0001<br>66 | 0.83918<br><.0001<br>66 | | | -0.31130<br>0.1069<br>28 | -0.07312<br>0.7116<br>28 | 0.22343<br>0.3578<br>19 | -0.18069<br>0.3482<br>29 |
| CT KCNH2 | -0.20507<br>0.2952<br>28 | 0.92163<br><.0001<br>66 | 0.86418<br><.0001<br>66 | | | -0.16477<br>0.4021<br>28 | -0.07597<br>0.7008<br>28 | 0.12778<br>0.6021<br>19 | -0.19862<br>0.3017<br>29 |
| CT CACNA1D | -0.01819<br>0.9884<br>3 | -0.50750<br>0.1631<br>9 | -0.44537<br>0.2296<br>9 | | | 0.99324<br>0.0741<br>3 | 0.99930<br>0.0238<br>3 | -1.00000<br>.<br>2 | -0.92248<br>0.2523<br>3 |
| CT CACNA1G | 1.00000<br>.<br>2 | -0.38033<br>0.4570<br>6 | -0.34388<br>0.5045<br>6 | | | 1.00000<br>.<br>2 | 1.00000<br>.<br>2 | .<br>.<br>1 | -1.00000<br>.<br>2 |
| CT KCNA5 | -0.45852<br>0.0185<br>26 | 0.66843<br><.0001<br>58 | 0.55315<br><.0001<br>58 | | | -0.44694<br>0.0221<br>26 | -0.47502<br>0.0142<br>26 | 0.33515<br>0.1885<br>17 | -0.17752<br>0.3757<br>27 |
| CT KCNJ2 | 0.02383<br>0.9061<br>27 | 0.91031<br><.0001<br>64 | 0.89763<br><.0001<br>64 | | | -0.20935<br>0.2946<br>27 | -0.11160<br>0.5794<br>27 | 0.35341<br>0.1502<br>18 | -0.13975<br>0.4781<br>28 |

| Pearson Correlation Coefficients<br>Prob > \|r\| under H0: Rho=0<br>Number of Observations | | | | | | | | | |
|---|---|---|---|---|---|---|---|---|---|
| | CT KCNJ4 | CT HCN4 | CT SCN5A | CT CACNA1C | CT KCNH2 | CT CACNA1D | CT CACNA1G | CT KCNA5 | CT KCNJ2 |
| Capacitance [pF] | | | | | | | | | |
| CT TNNT2 | | | | | | | | | |
| CT GAPDH | | | | | | | | | |
| Beat rate [bpm] | | | | | | | | | |
| Inter-beat interval [ms] | | | | | | | | | |
| Fast inward current amplitude [nA] | | | | | | | | | |
| Fast inward current density [nA/pF] | | | | | | | | | |
| Upstroke velocity [V/ms] | | | | | | | | | |
| APD90 [ms] | | | | | | | | | |
| CT KCNJ4 | | | | | | | | | |
| CT HCN4 | 0.69715<br><.0001<br>56 | | | | | | | | |
| CT SCN5A | 0.70826<br><.0001<br>60 | 0.83395<br><.0001<br>62 | | | | | | | |
| CT CACNA1C | 0.50424<br><.0001<br>60 | 0.78715<br><.0001<br>62 | 0.89932<br><.0001<br>66 | | | | | | |
| CT KCNH2 | 0.66709<br><.0001<br>60 | 0.79228<br><.0001<br>62 | 0.93961<br><.0001<br>66 | 0.82845<br><.0001<br>66 | | | | | |
| CT CACNA1D | -0.55434<br>0.1214<br>9 | -0.90516<br>0.0051<br>7 | -0.59789<br>0.0890<br>9 | -0.68836<br>0.0404<br>9 | -0.56775<br>0.1108<br>9 | | | | |
| CT CACNA1G | -0.19650<br>0.7090<br>6 | -0.51595<br>0.4841<br>4 | -0.29833<br>0.5658<br>6 | -0.36568<br>0.4759<br>6 | -0.28523<br>0.5838<br>6 | -1.00000<br>.<br>2 | | | |
| CT KCNA5 | 0.49670<br>0.0001<br>55 | 0.53913<br><.0001<br>54 | 0.69113<br><.0001<br>58 | 0.59693<br><.0001<br>58 | 0.72190<br><.0001<br>58 | -0.64581<br>0.0603<br>9 | -0.40892<br>0.4208<br>6 | | |
| CT KCNJ2 | 0.54435<br><.0001<br>59 | 0.77569<br><.0001<br>60 | 0.91569<br><.0001<br>64 | 0.88096<br><.0001<br>64 | 0.86678<br><.0001<br>64 | -0.62797<br>0.0702<br>9 | -0.16629<br>0.7529<br>6 | 0.53863<br><.0001<br>57 | |

**Supp. Table 6: Single-cell correlations for primary human atrial cardiomyocytes.** For every pairwise correlation, Pearson´s correlation coefficient r, the p-value and *n* is indicated. For expression parameters (CT), only positive cells are included in the analysis. Significant correlations ($p < 0.05$) are highlighted.

Pearson Correlation Coefficients
Prob > |r| under H0: Rho=0
Number of Observations

| | Capacitance [pF] | CT TNNT2 | CT GAPDH | Beat rate [bpm] | Inter-beat interval [ms] | Fast inward current amplitude [nA] | Fast inward current density [nA/pF] | Upstroke velocity [V/ms] | APD90 [ms] |
|---|---|---|---|---|---|---|---|---|---|
| Capacitance [pF] | | | | | | | | | |
| CT TNNT2 | | | | | | | | | |
| CT GAPDH | | 0.64314<br><.0001<br>40 | | | | | | | |
| Beat rate [bpm] | | | | | | | | | |
| Inter-beat interval [ms] | | | | | | | | | |
| Fast inward current amplitude [nA] | | | | | | | | | |
| Fast inward current density [nA/pF] | | | | | | | | | |
| Upstroke velocity [V/ms] | | | | | | | | | |
| APD90 [ms] | | | | | | | | | |
| CT KCNJ4 | | 0.71803<br><.0001<br>38 | 0.52165<br>0.0008<br>38 | | | | | | |
| CT HCN4 | | 0.65271<br><.0001<br>38 | 0.70499<br><.0001<br>38 | | | | | | |
| CT SCN5A | | 0.92851<br><.0001<br>40 | 0.65521<br><.0001<br>40 | | | | | | |
| CT CACNA1C | | 0.75201<br><.0001<br>39 | 0.79858<br><.0001<br>39 | | | | | | |
| CT KCNH2 | | 0.86837<br><.0001<br>40 | 0.66604<br><.0001<br>40 | | | | | | |
| CT CACNA1D | | 0.40383<br>0.0408<br>26 | 0.28054<br>0.1651<br>26 | | | | | | |
| CT CACNA1G | | 0.52913<br>0.0136<br>21 | 0.52760<br>0.0140<br>21 | | | | | | |
| CT KCNA5 | | 0.79660<br><.0001<br>40 | 0.71991<br><.0001<br>40 | | | | | | |
| CT KCNJ2 | | 0.52735<br>0.0019<br>32 | 0.65323<br><.0001<br>32 | | | | | | |

Pearson Correlation Coefficients
Prob > |r| under H0: Rho=0
Number of Observations

| | CT KCNJ4 | CT HCN4 | CT SCN5A | CT CACNA1C | CT KCNH2 | CT CACNA1D | CT CACNA1G | CT KCNA5 | CT KCNJ2 |
|---|---|---|---|---|---|---|---|---|---|
| Capacitance [pF] | | | | | | | | | |
| CT TNNT2 | | | | | | | | | |
| CT GAPDH | | | | | | | | | |
| Beat rate [bpm] | | | | | | | | | |
| Inter-beat interval [ms] | | | | | | | | | |
| Fast inward current amplitude [nA] | | | | | | | | | |
| Fast inward current density [nA/pF] | | | | | | | | | |
| Upstroke velocity [V/ms] | | | | | | | | | |
| APD90 [ms] | | | | | | | | | |
| CT KCNJ4 | | | | | | | | | |
| CT HCN4 | 0.43418<br>0.0081<br>36 | | | | | | | | |
| CT SCN5A | 0.77906<br><.0001<br>38 | 0.63107<br><.0001<br>38 | | | | | | | |
| CT CACNA1C | 0.74720<br><.0001<br>37 | 0.62625<br><.0001<br>37 | 0.82354<br><.0001<br>39 | | | | | | |
| CT KCNH2 | 0.72826<br><.0001<br>38 | 0.59691<br><.0001<br>38 | 0.87227<br><.0001<br>40 | 0.79846<br><.0001<br>39 | | | | | |
| CT CACNA1D | 0.24616<br>0.2356<br>25 | 0.22168<br>0.2869<br>25 | 0.28739<br>0.1546<br>26 | 0.36581<br>0.0721<br>25 | 0.28597<br>0.1567<br>26 | | | | |
| CT CACNA1G | 0.49896<br>0.0251<br>20 | 0.56495<br>0.0094<br>20 | 0.56293<br>0.0079<br>21 | 0.61373<br>0.0040<br>20 | 0.59517<br>0.0044<br>21 | 0.50094<br>0.0289<br>19 | | | |
| CT KCNA5 | 0.67437<br><.0001<br>38 | 0.60288<br><.0001<br>38 | 0.74573<br><.0001<br>40 | 0.80803<br><.0001<br>39 | 0.68701<br><.0001<br>40 | 0.44694<br>0.0221<br>26 | 0.58801<br>0.0051<br>21 | | |
| CT KCNJ2 | 0.37481<br>0.0345<br>32 | 0.38563<br>0.0322<br>31 | 0.57232<br>0.0006<br>32 | 0.61276<br>0.0002<br>31 | 0.56497<br>0.0008<br>32 | 0.30253<br>0.1712<br>22 | 0.78242<br>0.0001<br>18 | 0.58690<br>0.0004<br>32 | |
